# Assembling a true ‘Olympic Gel’ from >16,000 combinatorial DNA rings

**DOI:** 10.1101/2024.07.12.603212

**Authors:** Sarah K. Speed, Yu-Hsuan Peng, Azra Atabay, Krishna Gupta, Toni Müller, Carolin Fischer, Ilka M. Hermes, Jens-Uwe Sommer, Michael Lang, Elisha Krieg

**Affiliations:** Division of Polymer Biomaterials Science, Leibniz Institute of Polymer Research Dresden, Germany; Faculty of Chemistry and Food Chemistry, Technische Universität Dresden, Germany; Division of the Theory of Polymers, Leibniz Institute of Polymer Research Dresden, Germany; Institute for Theoretical Physics, Technische Universität Dresden, Germany; B CUBE - Center for Molecular Bioengineering, Technische Universität Dresden, Germany; Division of Physical Chemistry and Physics of Polymers, Leibniz Institute of Polymer Research Dresden, Germany; Cluster of Excellence Physics of Life, Technische Universität Dresden, Germany

**Keywords:** DNA nanotechnology, supramolecular chemistry, Olympic gels, programmable materials, soft matter engineering, simulations

## Abstract

Olympic gels are an elusive form of soft matter, comprising a three-dimensional network of mechanically interlocked cyclic molecules. In the absence of defined network junctions, the high conformational freedom of the molecules was previously theorized to confer unique mechanical properties to Olympic gels, such as non-linear elasticity and unconventional swelling characteristics. However, the synthesis of an Olympic gel exhibiting these intriguing features is challenging, since unintended crosslinking and polymerization processes are often favored over cyclization. Here, we report the successful assembly of a true Olympic gel from a library of DNA rings comprising more than 16,000 distinct molecules. Each of these rings contains a unique sequence domain that can be enzymatically activated to produce reactive termini that favor intramolecular cyclization. We characterized the genetic, mechanical, and structural characteristics of the material by next-generation sequencing, oscillatory rheology, large-scale computational simulations, atomic force microscopy, and cryogenic electron microscopy. Our results confirm the formation of a stable Olympic gel, which exhibits unique swelling behavior and an elastic response that is exclusively determined by entanglements, yet persists on long time scales. By combining concepts from polymer physics, synthetic biology, and DNA nanotechnology, this new material class provides a flexible experimental platform for future studies into the effects of network topology on macroscopic material properties and its function as a carrier of genetic information in biological and biomimetic systems. This work moreover demonstrates that exotic material properties can emerge in systems with a high compositional complexity that is more reminiscent of biological than synthetic matter.

## Introduction

Gels are versatile soft materials that exhibit unique mechanical and structural properties^1–3^. They consist of molecules that are crosslinked through either chemical bonds or physical (noncovalent) interactions. The resulting three-dimensional molecular network spans the entire volume of the gel, while the interstitial space is filled by a solvent. Gels enable a wide range of applications, ranging from drug delivery^4^, contact lenses^5^, and tissue engineering^6^ to ultrafiltration membranes^7^, autonomously controlled microfluidic channels^8^, and light harvesting scaffolds with stimuli-responsive properties^9^.

To date, the mechanical properties of gels are explained by relating the modulus (G) to the density of the junctions within the three-dimensional molecular network^10^. If these junctions were removed, the elastic material would turn into a fluid. In 1979, however, De Gennes conjectured^11^ that one could make “Olympic gels”—gels entirely devoid of network junctions but instead held together by the concatenation of cyclic molecules (Figure 1a). The lack of local junctions means that well-established concepts in polymer physics to link network structure to macroscopic properties cannot be easily applied to Olympic gels. Interestingly, in the absence of defined network junctions, overlapping rings can be displaced with respect to one another, lowering the entropic cost of macroscopic strain induced by external forces. Due to the topological nature of their crosslinks, Olympic gels are expected to have intriguingly different mechanical properties than classical gels^12^, in particular unique deformation^13^ and swelling^14^ characteristics.

The mechanical interlocking of macrocycles can also serve unique biological functions in nature. The unusual mitochondrial DNA of the microorganism *Kinetoplastea* (known as kinetoplast DNA, or kDNA) consists of several thousand concatenated DNA circles resembling a microscopic two-dimensional version of an Olympic gel^15,16^. Several members of *Kinetoplastea* are pathogenic parasites that cause dangerous diseases including Chagas disease, African sleeping sickness, and leishmaniasis. The kDNA network plays a critical role in the genetic function, post-transcriptional modifications, and cell division within *Kinetoplastea*, but how its topological structure might influence these biological functions is still under investigation^17,18^.

Despite the fascinating mechanical and functional properties of Olympic gels, their synthesis has proven remarkably challenging. True Olympic gels have to date only been demonstrated in computer simulations^14,19–21^. This is particularly surprising when considering that over the past 65 years diverse types of mechanically interlocked molecular rings (catenanes) have been created at the nanoscale^22–24^ and in 2D sheets^25^. Yet, the same approaches cannot be easily applied to create macroscopic materials; to form an Olympic gel one must achieve efficient cyclization of molecules at a very high concentration, as the newly formed macrocycles must overlap in space with other macrocycles to allow the formation of a sample-spanning concatenated network. However, at the required concentrations, a simple polymerization process is thermodynamically favored over cyclization for both reversible and irreversible reactions, such that the molecules are linked together to form linear chains^26^.

In recent decades, several ideas have been proposed to overcome the cyclization-vs-polymerization challenge^27,28^. However, computer simulations indicate that the required experimental conditions are often difficult to achieve in practice^19^. It has been reported that some degree of concatenation can be achieved in mixtures of cyclic polymers^29,30^. To date, no study has demonstrated the chemical synthesis of an Olympic gel and validated its distinct mechanical properties. A recent study suggested a synthetic route towards Olympic gels based on lipoic acid derivatives^31^. However, the proposed reaction mechanism implies the existence of classical covalent crosslinks, and distinct Olympic gel-like behavior was not reported. Two studies have reported a biologically-inspired approach for the synthesis of an Olympic gel using the help of the enzyme Topoisomerase II to concatenate plasmid DNA^32,33^. The enzyme transiently breaks and reforms plasmids, thus allowing previously separated DNA strands to pass through each other and thereby to interlock. This approach is elegant; however, there is strong evidence that the enzyme itself forms bridges between the DNA strands and thereby creates classical network junctions (as suggested by the latest study)^32,34^. The effect of these classical junctions dominates the mechanical characteristics of the material.

In the present study, we report a new approach for generating Olympic gels. This method is based on biotechnologically produced double-stranded DNA rings that are enzymatically processed to reversibly open and close at a predefined position. We have previously shown that the formation of intermolecular bonds in a hydrogel network can be promoted via the use of complex DNA crosslinker libraries containing up to 256 distinct recognition sites^35,36^. Here, we demonstrate that a similar approach can be used “in reverse” to favor intramolecular cyclization over intermolecular polymerization. By cloning combinatorial DNA sequence mixtures and amplifying them in *E. coli*, we generated a diverse library of DNA plasmids containing more than 16,000 distinct DNA sequences. These libraries were enzymatically processed, purified, and finally self-assembled to form an Olympic gel, as revealed by oscillatory rheology, atomic force microscopy, electron microscopy, and swelling experiments.

## Results and Discussion

### Concept and computer-assisted material design

To construct an Olympic gel, one needs a molecule that readily becomes mechanically interlocked but does not form classical junctions. We hypothesized that a polymer with two reactive termini would fulfill this requirement if the termini selectively recognize each other (analogous to a key and lock) but do not bind to other molecules in the vicinity. To achieve this highly selective cyclization while avoiding linear polymerization, a diverse mixture of components is needed, such that each of the molecules is likely to be surrounded only by molecules carrying different binding sites (Figure 1b). Assuming a large enough diversity of reactive termini, such mixtures can be expected to form an Olympic gel at high concentration^27^.

However, implementing this concept is not straightforward, as the set of orthogonal binding motifs available in organic or supramolecular chemistry is limited^37,38^. An exception is DNA, whose highly selective Watson-Crick-Franklin base pairing can be used to construct large libraries with orthogonal binding interactions^35,39^. We sought to insert such binding motifs into plasmids, naturally ring-shaped DNA molecules that are readily amplified in bacteria. To this end, we designed a synthetic DNA oligonucleotide termed the DLK (“Diversified Lock & Key”) insert for cloning into a plasmid backbone (Figure 1c). The DLK insert maximizes sequence complexity in the final plasmid library through the inclusion of a combinatorial *lock & key domain* consisting of 16 mixed nucleotides (N), where each N nucleotide can contain either of the four canonical bases A, T, C, or G with approximately 25% probability. This approach creates a mixture of 4^16^ (∼4 billion) unique lock & key domain sequences. The lock & key domain is flanked by two *nicking sites*, which are specific recognition sequences for the Nt.BbvCI nicking endonuclease. Enzymatic nicking on opposite sides of the strand creates a 20 nucleotide (nt)-long sticky end with the sequence GGN_16_CC, which can be reversibly opened and closed via thermal or chemical denaturation (Figure 1c). The DLK insert also includes two diversified *strand exchange protector* domains, each consisting of 9 N nucleotides. These domains flank the sticky ends and prevent spurious strand exchange that could otherwise take place at elevated temperatures due to strand fraying. Finally, the DLK insert contains two 26 nt-long *homology arms* that allow its integration into the plasmid via Gibson assembly^40^.

**Figure 1.**
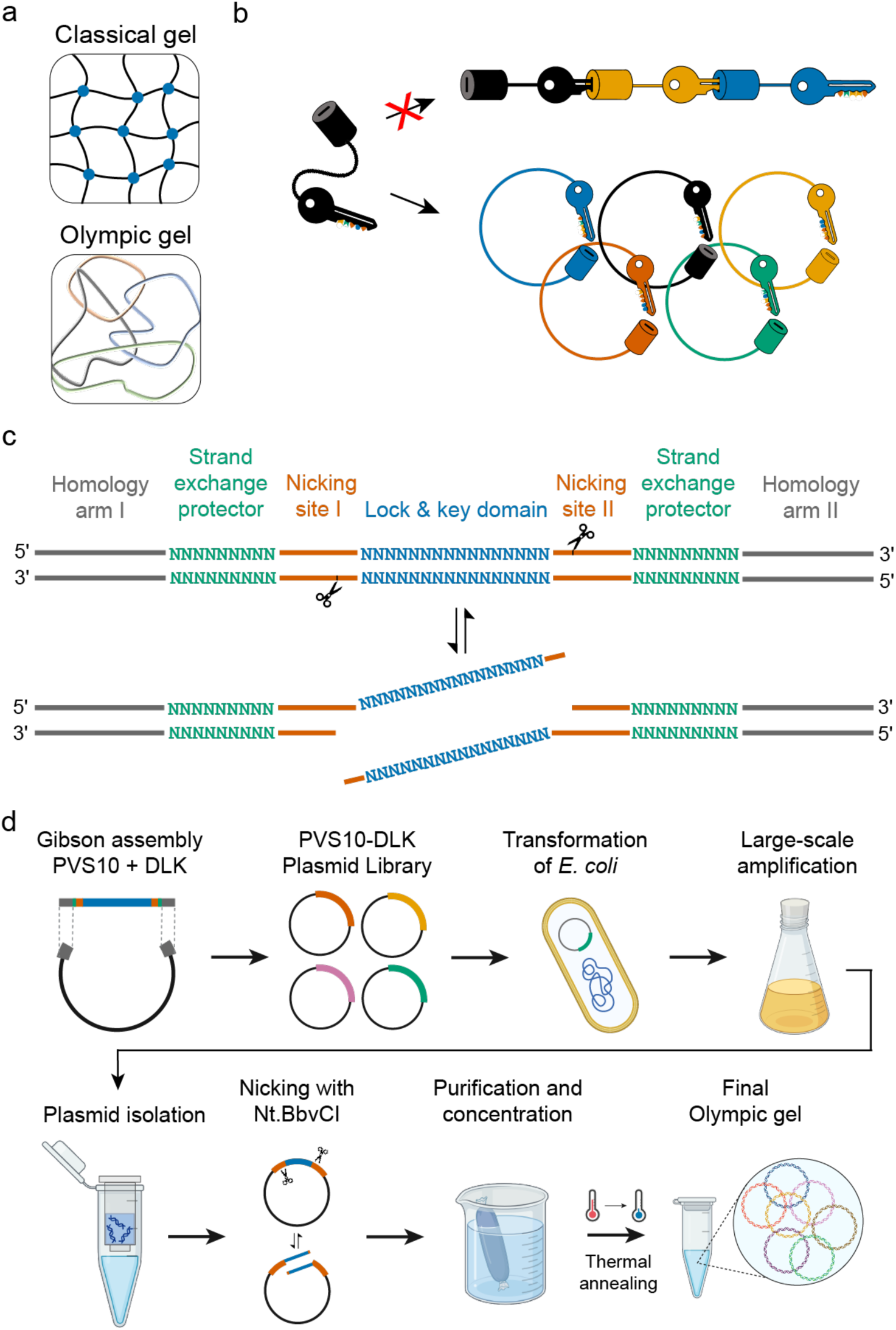
The synthesis of Olympic gels requires molecules that form concatenated rings instead of linear polymers or classical network junctions. **a)** Comparison between classical gels, which are crosslinked by either covalent or noncovalent junctions, and Olympic gels, which do not have distinct junctions but are held together by mechanically interlocked (concatenated) rings. **b)** Scheme of the key-in-lock model in which a hypothetical linear molecule with two reactive termini (a key and a lock) that bind to each other through selective recognition favors intramolecular over intermolecular binding. Given a library of molecules with a sufficient diversity of lock-and-key pairs, the polymerization of linear chains is suppressed, and the formation of an Olympic gel is thermodynamically favored. **c)** Design of the synthetic DLK insert for implementation of the key-in-lock model with DNA plasmid rings. Note that the scheme shows the insert in its double-stranded form, as present in the final plasmid product; Gibson assembly is conducted with the corresponding single-stranded oligonucleotide, and the complementary sequence is created *in situ* by the polymerase (see Methods). **d)** Scheme depicting the workflow in the biotechnological production of the Olympic gel.

Using nearest-neighbor thermodynamic calculations^41^, the 20 nt sticky ends were predicted to bind to each other with a minimum free energy (MFE) of −114 kJ mol^−1^ (see Methods). The melting temperature was predicted to be between 50–60°C. We therefore expected that at room temperature most plasmid rings are closed, but at low ionic strength or when annealed near the melting temperature, they would transiently open and allow for subsequent concatenation. To further guide our experimental design, we used bond-fluctuation model (BFM) computer simulation data^42^ to predict the critical gelation concentration (*c_gel_*) and the average number of expected concatenations per ring (propensity for concatentation, *f_n_*) at different concentrations (Supplementary Section S2, Supplementary Figure S5). These simulations show that the structure of Olympic gels made of monodisperse rings depends solely on *_fn_* ∝ *Nc* for rings with ideal conformations, where *N* is the degree of polymerization (i.e., the size of the rings), and *c* is the mass concentration of the rings. Thus, the propensity of concatenation can be systematically varied by either changing the size or the concentration of the rings. Based upon the simulation data, *c_ge_*_l_ was predicted as 0.11 wt%. This point corresponds to a universal overlap number (*P*) of *P* = 0.36 for monodisperse rings with random walk conformations, above which a cyclic polymer can form an Olympic gel (see Supplementary Section S2 for details). For ideal plasmid solutions of constant molar mass, such as for our series of gels, *f_n_* can be estimated as *f_n_* ≈ *c/c_gel_* ^19^. The linear growth of *f_n_* with the concentration was also found in simulations of kDNA minicircle networks^43^. Based on these calculations, we prepared plasmid solutions with weight fractions ranging from approximately one order of magnitude below to one order of magnitude above the predicted value for *c_gel_* (0.01–0.8 wt%).

### Biotechnological production of the Olympic gel

We applied classical molecular biology techniques for the production of the diverse plasmid library (Figure 1d, see Methods section). We first integrated the synthetic DLK insert into a PVS10 plasmid backbone^44^ using Gibson assembly^40^ to produce the initial PVS10-DLK plasmid library (Supplementary Data 1). This library was amplified on a large scale in *E. coli* and subsequently purified, yielding several milligrams of plasmid DNA per synthesis batch. A portion of this purified plasmid product was enzymatically nicked at the two predetermined locations using Nt.BbvCI nicking endonuclease. This step removed any supercoiling and activated the DLK insert to allow for reversible ring opening and concatenation (Figure 1d). The double-nicked PVS10-DLK plasmids are hereafter referred to as “Olympic” samples. To provide a control, we digested a second portion of the purified plasmid product with a different nicking enzyme, Nb.BsmI. Nb.BsmI nicks the plasmid in only one location, remote from the DLK insert. Introducing this nick removes supercoiling and unwinds the plasmid into a conformationally relaxed circular form. These circles are similar to the double-nicked plasmids, but lack the ability to open and close reversibly. The Nb.BsmI-treated samples served as the primary control to demonstrate the impact of the concatenated structure of the Olympic gels. We verified the nicking efficiency in both the Olympic and relaxed control samples through an analytical HindIII restriction enzyme digestion followed by denaturing agarose gel electrophoresis (AGE) analysis (Figure 2a). The expected digest patterns were observed for both the Olympic and control samples, confirming the successful and near-quantitative nicking of the plasmid libraries.

To explore the formation of Olympic gels, we prepared different concentrations of the plasmid samples (0.01–0.8 wt%) in aqueous solution containing 50 mM sodium chloride. High salt concentration and divalent cations were avoided, as these can promote nonspecific interactions between DNA strands^45^. The samples were annealed at 50 °C to trigger ring opening and to allow the rings to pass through one another, followed by incubation at room temperature to allow concatenation. All control samples were treated identically.

To verify concatenation of the plasmid rings, we first analyzed solutions of the Olympic and control samples by native AGE (Figure 2b, Supplementary Figure S6, and Supplementary Section S3). Olympic samples contained both relaxed circular (80%) and linear (20%) species. The linear form was expected, since pipetting-induced shear forces are strong enough^46^ to dissociate the lock & key domain during sample loading. As per their design, control plasmids were present in relaxed circular form, with small amounts of supercoiled circular plasmids but no linear form detectable. Importantly, as the Olympic samples approach the overlap concentration (0.11 wt%), a large fraction of DNA is trapped near the well, while control samples remain largely dissociated. The intensity of this band in relation to the free circular plasmid band increases with increasing concentration, which is indicative of concatenation of the plasmid rings leading to the formation of larger clusters of rings in solution (Figure 2b). Unlike the control samples, Olympic samples above 0.1 wt% were gel-like and difficult to pipet and dilute, leading to notable artifacts in AGE (Supplementary Figure S6). Surprisingly, AGE did not resolve clear bands for small interlocked multimers, despite these being observed in AFM (Figure 3c,d, Supplementary Figure S12). We suspect that transient ring opening can take place as small plasmid clusters migrate through the agarose gel for several hours, which would lead to irreversible ring separation and diminishing multimer bands. Overall, AGE demonstrates that plasmid ring concatenation requires the fulfillment of two criteria: 1) the ability of the plasmids to reversibly open and close and 2) a concentration approaching the predicted *c*_gel_ value that would allow overlap of the rings in solution.

### Sequencing reveals highly diversified lock & key domains with orthogonal binding

Achieving a high number of unique sequences within the DLK insert of the final plasmid library is key for generating an Olympic gel^19^. We speculated that large-scale plasmid amplification in *E. coli* could, in principle, lead to distortions in the prevalence of distinct sequences, as some sequences might be preferentially amplified. In order to verify that the plasmid library sequences were still widely distributed after amplification, we characterized the DLK insert in both the original Gibson product before transformation and the final purified PVS10-DLK library (obtained from the mixing of five parallel transformations) by next-generation sequencing (Figure 2c–e, Supplementary Data 2).

We obtained and analyzed more than 100,000 sequences of properly merged paired end reads for both the Gibson product and the final library. Using a custom Python script (Supplementary Data 2), we identified instances of Nt.BbvCI nicking sites in order to extract sequences of interest. We found that ∼98.5% of the analyzed sequences contained proper nicking sites, both in the initial Gibson product as well as the final library (Figure 2d), demonstrating that DLK insertion into the plasmid was efficient and that almost all of the DNA rings can be activated by nicking. Importantly, the lock & key domain as well as the two strand exchange protector domains exhibited high sequence diversity in the final library (Figure 2c,d). A slight bias was found in favor of adenine and thymine (58%) over cytosine and guanine (42%). This bias existed already in the initial Gibson product (Supplementary Figure S11a), reflecting a known bias in the phosphoramidite-based synthesis of the commercial oligonucleotide^47^. The amplification in *E. Coli* did not introduce additional biases for specific bases.

We then computed the total number and prevalence of unique sequence variants within the DLK insert (Figure 2d,e). The pool of 140,543 identified sequences in the initial Gibson product contained 125,815 unique sequences (89.5%), demonstrating that the full library’s complexity substantially exceeded the number of total sequence reads, as predicted. After large-scale plasmid amplification, we identified 16,307 unique variants in the final library, where the most abundant sequence had a prevalence of 0.067%. This decrease in unique sequences was also expected, since the number of distinct plasmid sequences cannot exceed the initial number of transformed bacterial cells. The final library had a Simpson Diversity Index (S) of 1.6 x 10^−4^, and a Shannon Information Entropy (H) of 13 bit, underscoring an exceptional compositional complexity in the realm of artificial materials. For concentrations exceeding the overlap concentration by a factor of 10, all plasmids must have prevalences significantly below 10% to suppress their polymerization^26^ and to allow Olympic gelation^19^. Thus, the experimentally verified maximum sequence prevalence is more than two orders of magnitude lower than required, such that each plasmid has a negligible chance of binding a complementary lock & key domain of another plasmid it overlaps with.

To confirm the high selectivity of individual variants for their reverse complements on the same plasmid backbone, we conducted nearest-neighbor thermodynamic predictions on the lock & key domains for all ∼266 million pairwise interactions between the 16,307 unique sequences in the final library (Supplementary Data 3). The calculated interactions between the 10 most abundant lock & key domains in the final library are shown in Figure 2f. The variants have a high predicted affinity towards their reverse complements, with MFE_specific_ = −106.9 ± 10.6 kJ mol^−1^, which is in good agreement with the initially predicted value of −114 kJ mol^−1^. In contrast, the non-complementary pairs averaged a MFE_nonspecific_ of −25.4 ± 6.5 kJ mol^−1^ (Figure 2g). The large difference (Λ1MFE ∼ 81.5 kJ mol^−1^) confirms that the intramolecular pairing of lock & key domains is highly selective and the final PVS10-DLK plasmid library exhibits near-ideal orthogonal binding under thermodynamic equilibrium conditions.

**Figure 2.**
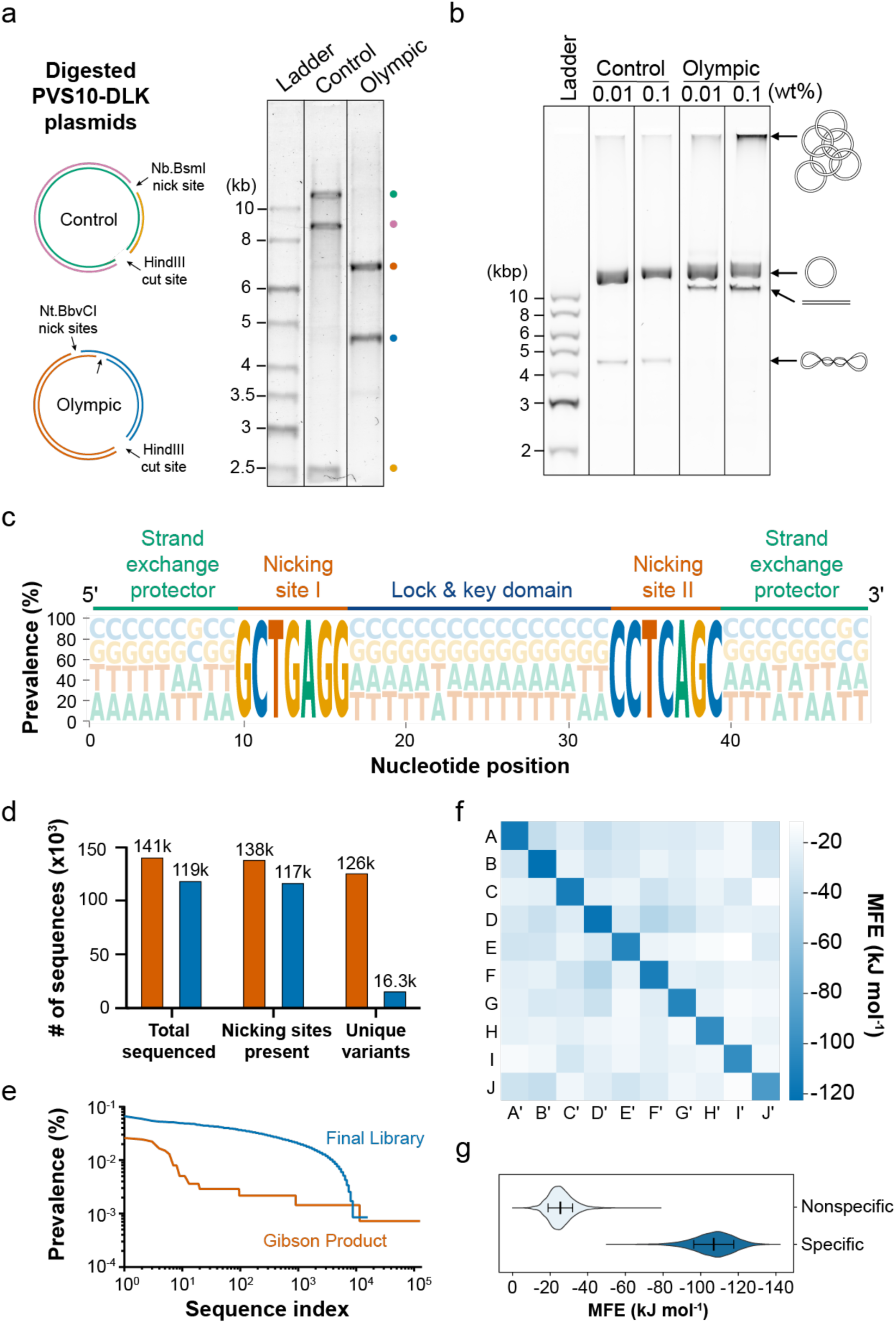
The PVS10-DLK plasmid library is efficiently activated by the nicking enzyme, while the high sequence diversity of the DLK insert promotes selective ring closure and concatenation. **a)** Alkaline denaturing AGE of Olympic plasmids and control plasmids used in this study. Prior to the experiment, the plasmids were digested by HindIII restriction enzymes to generate linear products to allow verification of the expected fragment lengths (kb = length of single-stranded DNA in kilobases). **b)** Native AGE of Olympic plasmids and control plasmids that were annealed at different concentrations (kbp = length of double-stranded DNA in kilobase pairs). The full gel showing a wider range of concentrations is shown in Supplementary Figure S6. **c)** Sequence logo showing the prevalence of the four bases within the DLK insert in the final PVS10-DLK library. The size of each letter corresponds to the probability of the indicated nucleotide to appear at that position. **d)** Next-generation sequencing data for the Gibson product prior to its transformation into *E. coli* (orange) and the final purified PVS10-DLK plasmid library (blue). **e)** Comparison of the prevalence between sequence variants (with nicking sites present) in descending order. **f)** Heatmap showing the predicted minimum free energies (MFE) for the interactions within the ten most prevalent lock & key domains (A, B, …, J) with the general sequence GGN16CC and their reverse complements (A’, B’, …, J’). A larger matrix containing all ∼266 million predicted hybridization interactions within the full library of 16,307 unique DLK sequences is shown in Supplementary Data 3. **g)** Violin plot showing the distribution of MFE values for specific sticky end interactions (corresponding to diagonal matrix entries; n = 16,307) and nonspecific interactions (corresponding to off-diagonal entries; n = 265,918,249).

### Structural characterization reveals an interconnected network of plasmid rings

To better understand the structural characteristics of the Olympic gel, we performed atomic force microscopy (AFM) imaging for plasmid samples at concentrations below *c_gel_* and cryogenic scanning electron microscopy (cryo-SEM) imaging at concentrations above *c_gel_*. Below *c_gel_*, all PVS10-DLK plasmid solutions are fluids, independent of enzymatic processing. Above *c_gel_*, Olympic plasmids become gel-like, while the original (non-nicked) plasmids and the relaxed controls remain as fluids. The transition to a gel-like material was dependent on thermal annealing, which is in agreement with the requirement for enzymatically activated plasmids to be opened and closed in order to allow efficient concatenation (Figure 3a).

We expected that annealing Olympic plasmids at concentrations just below *c_gel_* would yield small clusters of concatenated plasmids that could be imaged by AFM. The annealed samples were diluted to 5 ng μL^−1^ immediately prior to imaging to ensure that multiple plasmids were unlikely to overlap on the surface, unless physically linked together. AFM images revealed circular (closed) plasmids, linear (open) plasmids, and larger plasmid clusters (Figure 3c, Supplementary Figure S12). We measured the approximate contour length of each continuous surface object with a semi-automated script, yielding a weight-average length distribution (Figure 3b, see Methods). A single PVS10-DLK plasmid has an expected contour length of approximately 3.9 µm (11,344 base pairs × 0.34 nm per base pair in a DNA duplex). Olympic samples showed distinct peaks for the monomeric as well as multimeric structures (Figure 3b). These multimeric clusters were absent in the control and the original (non-nicked) plasmids. In some images, the mechanical interlocking of the Olympic plasmids could be visualized (Figure 3d). We did not observe linear multimers in any of the Olympic plasmid AFM images (n = 39, Supplementary Figure S12), confirming that undesirable linear polymerization is effectively suppressed. Overall, AFM imaging demonstrates that annealed Olympic samples below *c_gel_* contain linear (open) monomers, circular (closed) monomers, as well as mechanically interlocked multimers.

We further aimed to characterize the microscopic structure of the network at high concentration (above *c_gel_*) where the formation of a macroscopic gel was observed. Imaging 3D molecular networks in aqueous media is challenging, since typical sample preparation often causes prominent artifacts, such as network collapse due to drying or ice crystal formation due to freezing. We therefore characterized the sample by cryo-SEM, a technique that reduces preparative artifacts by transforming and maintaining the specimen in a vitrified state^48^. To this end, an Olympic gel sample was prepared at 0.25 wt%—approximately twice the predicted *c_ge_*_l_ value. The annealed product was subsequently frozen at high pressure to maintain the hydrated state and to conserve fine structural features within the gel (see Methods).

During imaging, the sample was subjected to sublimation of water at −90 °C to reveal the material’s internal structure^49^. The images show a mesh-like, interconnected network that is akin to network structures observed in conventional hydrogels^9,48^ (Figure 3e, Supplementary Figure S13). The prolonged sublimation at −90 °C caused local deformations that created pores within the gel network. This effect exposed fine strands of material that can be seen to occasionally span across gaps in the gel (Figure 3e, insets 1 and 2), which we attribute to plasmids or plasmid bundles that are stretched out as the vitrified water slowly retracts during sublimation. The observed interconnected structure with interstitial cavities supports the notion that the plasmid strands form mechanically stable connections within the gel network. Cryo-SEM imaging of the control sample and non-nicked plasmid sample that were treated identically do not show any network structure (Supplementary Figure S13, S14), further corroborating the notion that ring opening and closing is required to form a stable network structure.

**Figure 3.**
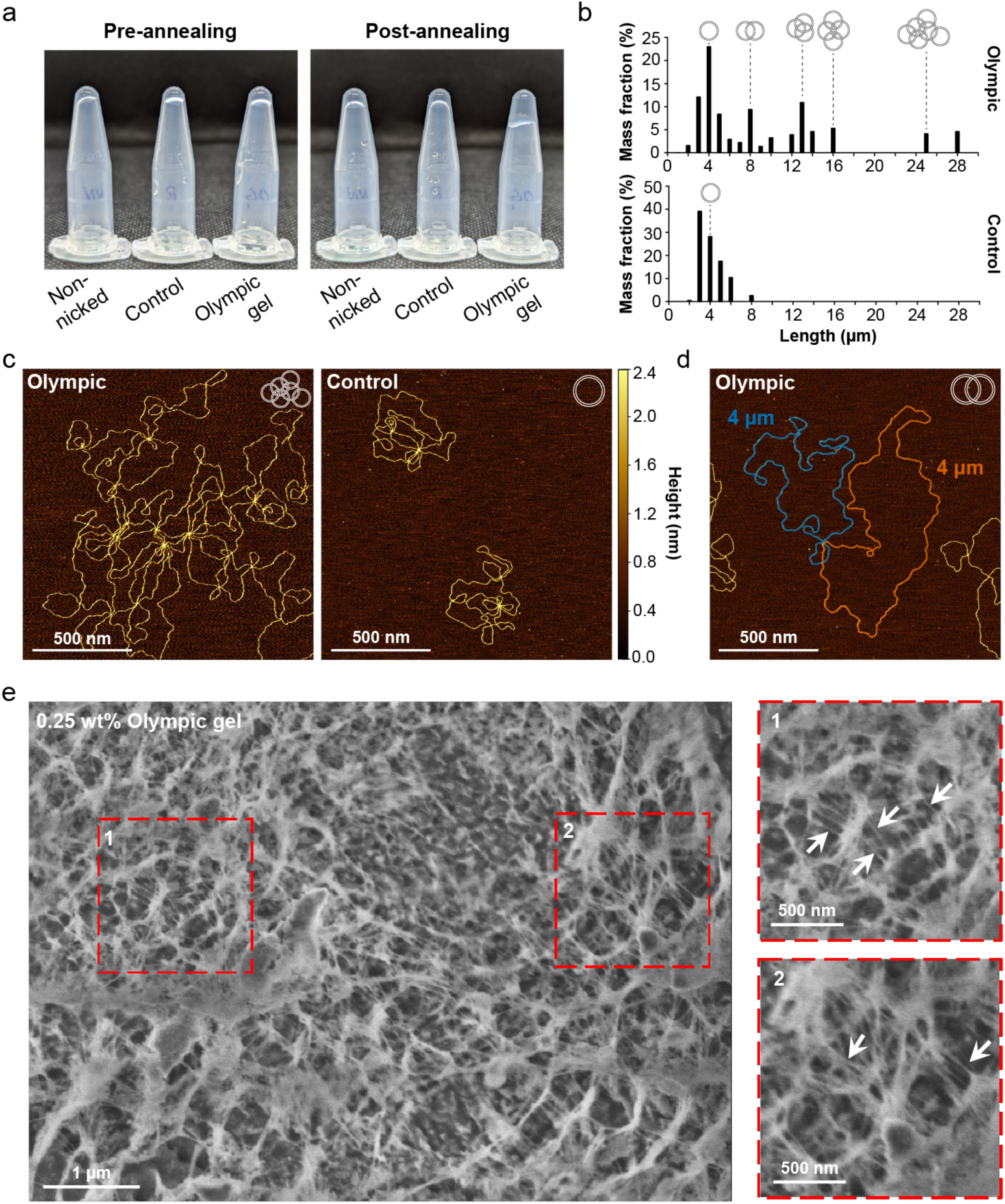
Structural characterization reveals the interconnected molecular network of plasmid rings constituting the Olympic gel. **a)** Tube inversion test for non-nicked, control, and Olympic gel samples at 0.125 wt% before and after annealing. **b)** Mass fraction of DNA present for different contour lengths of continuous surface objects in the AFM images of Olympic and control samples (n = 103 for Olympic; n = 74 for the control). **c)** AFM images of Olympic and control plasmids after annealing slightly below *cgel*. Additional representative images are shown in Supplementary Figure S12. **d)** AFM image of two mechanically interlocked Olympic plasmids with their corresponding lengths. The plasmids were colored blue and orange for clarity. The original uncolored image is shown in Supplementary Figure S12. **e)** Cryo-SEM image of the 0.25 wt% Olympic gel. Arrows in insets indicate suspected plasmids or plasmid bundles that are stretched out due to mechanical deformation of the network in response to sublimation. Additional cryo-SEM images of control plasmids and non-nicked plasmids are shown in Supplementary Figures S13 and S14.

### Computer simulations predict trends in mechanical and physical properties

To understand the mechanical properties of the concatenated networks, we carried out large-scale computer simulations in which randomly concatenated polymer rings with monodisperse size were sheared at varying applied stress, *σ* (Figure 4a, Supplementary Section S1). For optimal performance, the simulations were carried out for flexible polymers while varying the degree of polymerization. This yields in effect a variation of the overlap number, *P,* and the average number of concatenations per ring, *f*_n_, over a similar range as in the experimentally measured concentrations.

In the simulations, the strain (*γ*) was computed for different applied stresses until an equilibrium state was reached (Supplementary Figures S1, S3). The resulting stress-strain data of the strongly concatenated samples (Figure 4b) reveals a sub-linear stress-strain relation *σ* ∝ *γ*^0.78±0.03^ over a broad range of strains 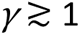. The observed exponent falls in between the linear dependence expected for entanglement-free systems and a constant asymptotics at large strains expected as limiting case for the slip-tube model in uni-axial deformation^50^. At the lowest applied forces, we observe an increasing strain (indicating an increased “slip length”) for increasing degree of polymerization of the rings. This behavior crosses over to a universal stress-strain relation at strains in the order of 100% or larger.

The increased strain for Olympic gels at low applied forces had been previously predicted^13^, however, questions remain regarding the cross-over towards larger applied stress. A larger slip length with increasing *N* agrees qualitatively with the predictions of the slip link model for rubber elasticity^51,52^. It is difficult to extract such a dependence from stress-strain data of conventional networks due to the additional contribution from the cross-links. We anticipate that our deformation data contributes valuable information on the deformation behavior of entangled networks and may help to improve existing models for rubber elasticity^50^. Finally, we simulated the deformation of the less concatenated samples with *N* = 128 and *N* = 64. For *N* = 128 (Supplementary Figure S2), a clearly lower modulus with a linear stress-strain relation was observed (see Supplementary Section S1), whereas equilibrium strain was not reached for the *N* = 64 sample within the available simulation time. Altogether, the present set of simulation data clarifies that rheological data at low strains for the samples with a large number of concatenations deserve particular attention.

**Figure 4.**
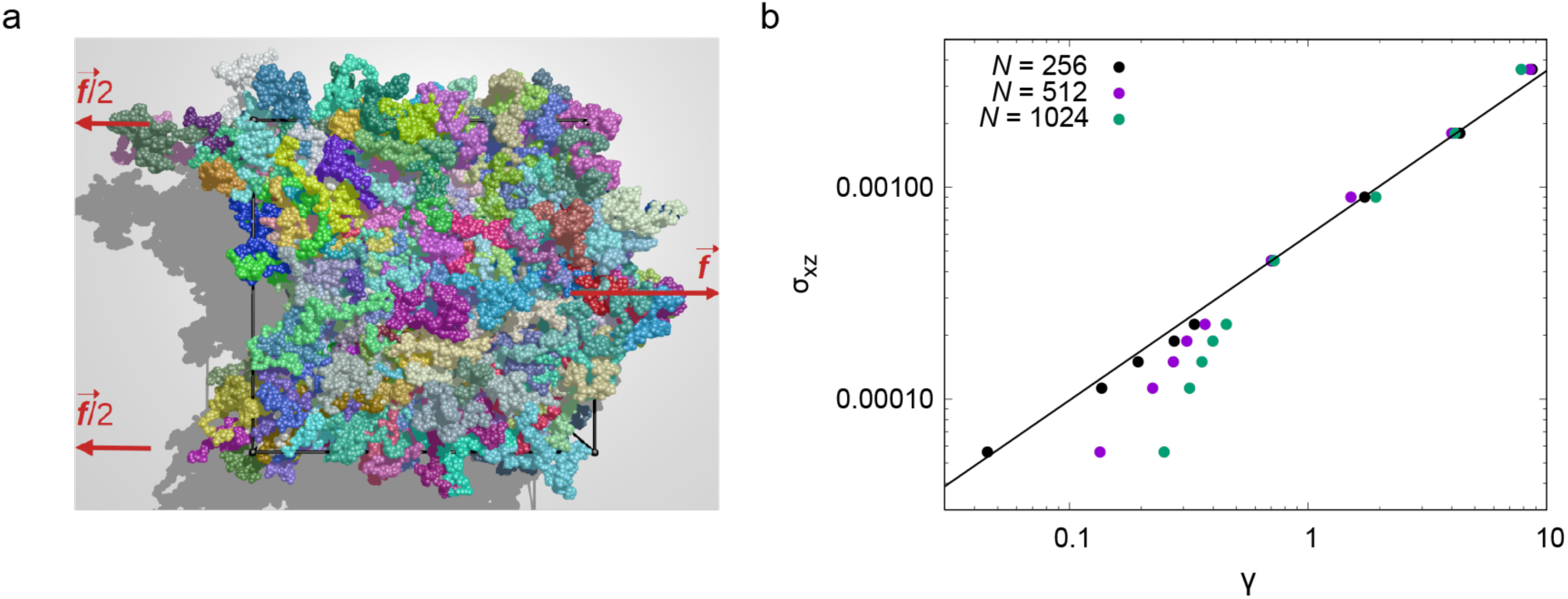
Computer simulations predict physical properties of the Olympic gel under different mechanical strains and varying degrees of concatenations. **a)** Example for a sheared Olympic gel with degree of polymerization (*N*) of 1024, a force f = 0.0036 kBT per unit length, and an average shear of 0.72. The applied force pulls the middle of the sample to the right and the top and bottom to the left. Each ring has a random color. Rings are displayed with respect to the center of mass position at the beginning of the shear considering periodic boundary conditions. Simulation snapshots for other values of *N* are shown in Supplementary Figure S4. **b)** Simulation results of ideal concatenated rings under shear stress σxz (in kBT per unit volume) for *N* = 256, 512, 1024. Increasing *N* leads to increasing ring overlap and increasing propensity for concatenations. The line indicates the relationship *σ* ∝ *γ*^0.78^.

### The Olympic gel exhibits unique swelling characteristics

To experimentally characterize macroscopic material properties, we first compared the swelling and dissolution behavior of the annealed Olympic gel and control samples at different concentrations. The samples were stained with a DNA-binding dye, covered with an excess of buffer solution, and subsequently imaged in a confocal microscope (Figure 5a,b). All control samples dispersed within 48 hours, even at the highest concentration (0.8 wt%), demonstrating that the rings neither crosslink nor entangle permanently (Supplementary Video 1). The same was observed for non-nicked plasmids (Supplementary Video 2). The time scale of the dispersion of the control plasmids is in good agreement with their expected rate of diffusion (Supplementary Section S6). In contrast, the Olympic gel samples exhibited a strongly concentration-dependent behavior: at high concentrations with *c* > *c_gel_* they swell to a maximal volume and subsequently retain their swollen shape (Supplementary Video 3). A slow shrinkage after the initial rapid swelling is observed for all Olympic gel samples. This is likely caused by diffusion of non-concatenated (e.g., leftover linear open) DNA out from the gels (Supplementary Section S6). The end-point volume was imaged after swelling for one week (Figure 5c). The Olympic gels at concentrations above the predicted *c_gel_* (0.125–0.8 wt%) remained intact. In contrast, Olympic plasmids below this threshold (0.025 wt% and 0.05 wt%) dispersed over time, similar to the control plasmids. This finding demonstrates that the Olympic gel forms a permanently connected network only above the theoretically predicted value of *c*_gel_.

The concentration-dependent trend of the volume change further corroborates the topological nature of the material. Classically crosslinked polymer gels generally swell to larger volumes when prepared at a higher polymer content^53^. In contrast, for an entangled polymer network, the largest swelling ratio is expected just above the gelation concentration, which then decays and approaches a constant swelling ratio independent of concentration (Supplementary Section S5)^54^. Figure 5c shows the experimentally measured end-point swelling ratio (after a one-week incubation), which is distinctly different from classically crosslinked gels but in good qualitative agreement with the expectations for Olympic gels. This provides strong evidence that the elastic properties of the gels on long time scales are dominated by permanent entanglements between the cyclic DNA strands.

### Rheology validates the predicted trends for Olympic gel viscoelasticity

To study the mechanical properties of the gel, a set of samples was prepared above the predicted *c_gel_* (0.125–0.8 wt%) and studied by a broad set of rheological experiments. The systematic comparison between Olympic gel and control samples aimed to isolate the effect of *permanent* mechanical bonds from *transient* plasmid entanglements. The samples were first subjected to step-strain experiments to study their stress-relaxation behavior (Figure 5d). Control plasmids showed a logarithmic decay, reaching 95% completion within 1000 seconds for the highest concentration (0.8 wt%). Notably, this decay was slower than the stress-relaxation of the original non-nicked plasmids (Supplementary Figure S10). The Olympic gel samples showed a more complex relaxation profile: a similar decay to the control samples was observed for the first few seconds, but an additional much slower decay contribution is present that is characteristic for a gel converging to a non-zero storage modulus (Figure 5d). This confirms that in the Olympic gel a substantial part of the deformation energy is stored elastically over long timescales.

In all Olympic gel samples, the storage modulus (*G*ʹ) exceeded the loss modulus (*G*ʹʹ) over a wide range of frequencies, confirming that stable and consistent gels were formed (Figure 5g). The loss factors at 0.001 Hz ranged from 0.5 (for 0.125 wt%) to 0.1 (for 0.8 wt%) (Figure 5e). A shallow peak in the *G*ʹʹ data was observed at ∼0.1 Hz for the highest concentration, 0.8 wt%. This peak likely corresponds to the relaxation of residual linear chains, that is, of rings that have not closed (see Supplementary Sections S3, S4). In mixtures of cyclic and linear polymers of the same molar mass, this particular relaxation is the slowest process^55,56^. The *G*ʹ data at the lowest measured frequency (0.001 Hz) matches the expected concentration scaling of semi-dilute entangled solutions (*G*ʹ ∝ *c*^2.3^) (Figure 5e) and not the scaling for conventional gels (*G*ʹ ∝ *c*)^53^. A quantitative analysis based upon the slip-tube model of rubber elasticity provides an alternate estimate for *f*_n_, which exceeds our original estimate using computer simulation data and plasmid conformations by only 13% (see Supplementary Section S4.D for details). Therefore, our estimates for *f_n_* based on simulations and rheology data are consistent, demonstrating that the elastic behavior of the Olympic gel samples is dominated by entanglements.

**Figure 5.**
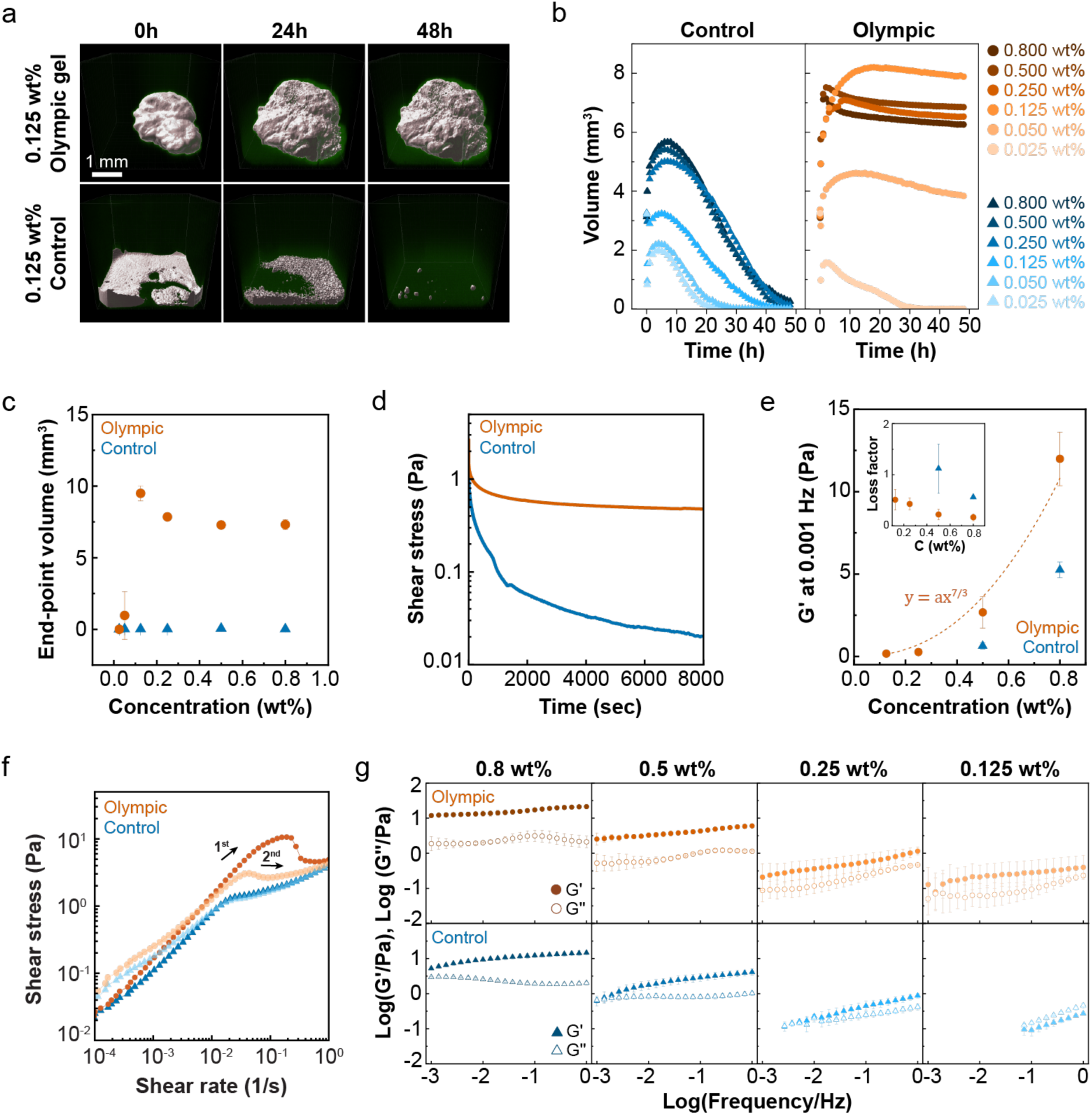
Rheology and confocal microscopy demonstrate the mechanical effects of plasmid concatenation and the unconventional swelling characteristics of the Olympic gel. **a)** Confocal images of the 0.125 wt% Olympic gel sample swelling in buffer (top row) and dissolution of the 0.125 wt% control sample under identical concentrations and buffer conditions (bottom row). The gel volume is rendered with a white surface; DNA-binding dye is shown in green. The white scale bar applies to all images. **b)** Swelling of samples immersed in Tris-NaCl buffer for 48 hours and **c)** Final gel volume after swelling for one week. Data are shown as the mean ± s.d. (n = 3 physical replicates). **d)** Stress-relaxation curves of 0.8 wt% samples at 20 °C. The shear stress was recorded over time at a fixed strain of 15%. **e)** Plot of the storage modulus, *Gʹ*, measured at 1% strain and 0.001 Hz in the frequency sweeps. Data are shown as the mean ± s.d. (n = 2 physical replicates, each averaged from two separate measurements). The dashed line and equation correspond to the analytical expression for the entanglement model. Factor *a* was fitted with a value of 18.2. The inset shows measured loss factors at 0.001 Hz. **f)** Flow curves of Olympic gel and control plasmids at 0.5 wt%. For each sample, two consecutive repeat experiments were carried out. The Olympic gel exhibits a yield point at ∼400% strain (∼0.2 s^−1^ shear rate). Flow curves for a range of concentrations are shown in Supplementary Figure S7. **g)** Frequency sweeps at 1% strain at 20 °C after annealing. Data are shown as the mean ± s.d. (n = 2 physical replicates, each averaged from two separate measurements).

In contrast to the Olympic gel, the *Gʹ* curves of control plasmids exhibited faster decay with decreasing frequency and larger loss factors (Figure 5e,g). Measurements at 0.125 wt% and 0.25 wt% concentrations approached the torque limit of the instrument, preventing the recording of their moduli in the low frequency range. Unlike in Olympic samples, control plasmids displayed a trend for a crossover between *Gʹ* and *Gʹʹ* (indicating the transition to a viscous liquid), which shifted to lower frequencies with increasing concentration (Figure 5g). These differences highlight the dominant contribution of long-lasting concatenations to the Olympic gel’s elasticity. Even though *Gʹ* values of control plasmids were consistently lower than for the Olympic gel (Figure 5e), they were surprisingly high even at low frequencies, in particular at the highest tested concentration. Such unexpectedly slow relaxation dynamics in plasmid solutions were reported in multiple recent studies, suggesting the involvement of unusually slow processes that are yet to be fully understood^57–60^.

In addition to oscillatory and step-strain experiments, we also subjected the material to rotational rheology. Flow curves show that Olympic gels exhibit substantially higher resistance to deformation than control plasmids (Figure 5f, Supplementary Figure S7). Distinct yield points are observed in all Olympic gels, exceeding yield stresses measured in control plasmids for all concentrations. When subjected to shear beyond the yield point, the Olympic gels break down. Measuring a second flow curve immediately after breaking the material in the first cycle results in a strongly decreased resistance to deformation. The observed hysteresis indicates that ring closure for renewed plasmid concatenation requires more than one minute (the delay between the flow experiments), which is consistent with the presence of open PVS10-DLK plasmid species in AGE and AFM experiments. Overall, rotational rheology suggests that the large deformation resistance of the Olympic gel samples under flow stems from mechanical bonds, distinctly different from the transient plasmid entanglements in control samples.

The strain amplitude at the yield point in flow curves coincided with the crossover of *Gʹ* and *Gʹʹ* in amplitude sweeps (e.g. ∼400% at 0.8 wt%; Supplementary Figure S8). This crossover shifted to larger strain values with decreasing concentration in Olympic gels (0.25– 0.8 wt%), indicating reduced plasmid mobility and tighter network formation at higher concentrations. The sub-linear increase of shear stress versus applied strain is a clear indication that elasticity is dominated by permanent entanglements. Quantitatively, the measured exponents are slightly below the exponent of 0.78 measured in the simulations (Supplementary Figure S9; Supplementary Section S4.E). Based upon the flow curves, we expect that an increasing portion of the rings is forced to open for high increasing strains explaining this small difference. The hysteresis in the amplitude sweeps of Olympic gel samples confirms incomplete network recovery and slow reformation of closed rings between cycles (Supplementary Figure S8). In contrast, the control plasmids in the same concentration range showed crossover points that were largely independent of concentration and remained around ∼180% strain. The control samples did not show hysteresis between the first and second sweeps, which is expected for entangled polymer solutions near the equilibrium.

Overall, the rheological data support the formation of a stable Olympic gel from double-nicked PVS10-DLK plasmids at concentrations ≥ 0.125 wt%, very close to the theoretically predicted *c_gel_* value of 0.11 wt%. Carefully controlling for temporary entanglements and weak plasmid interactions proved critical in this study, since gel-like behavior on short and intermediate timescales is also observed for conformationally relaxed control plasmids at high concentrations. The distinct characteristics of the concatenated structure of the Olympic gel versus temporary entanglements in relaxed control plasmid DNA become particularly dominant on long timescales, as manifested in swelling experiments (Figure 5a–c), stress-relaxation (Figure 5d), and flow experiments (Supplementary Figure S7).

## Conclusions

In the present study, we have combined the tools of synthetic biology, DNA nanotechnology, computer simulations, and theoretical polymer physics to synthesize and elucidate the properties of an Olympic gel—a new form of soft matter with unique properties. The key innovation is the use of highly diversified DNA recognition sites (“lock & key domains”) that were integrated into plasmid backbones and activated through enzymatic nicking. The lock & key domains serve as reactive termini that cause activated plasmids to favor self-cyclization over polymerization, even at high concentrations. In contrast to previous studies, this strategy allows efficient purification of the material and subsequent gelation while precluding spurious formation of classical crosslinks. This approach allowed an accurate characterization and investigation into the physical properties of Olympic gels.

Our combinatorial approach represents a rare example of how unique physical properties can emerge in materials comprising a large diversity of components. Notably, high compositional complexity has been previously shown to yield fascinating emergent properties in *high-entropy alloys*^61^—inorganic materials containing five or more different metals, whose “*cocktail effect*”^62^ provides a vast design space and exotic mechanical characteristics. We expect that other interesting material architectures may be achieved by using combinatorial synthetic biology in soft matter engineering.

Through microscopic imaging (AFM, cryo-SEM, and confocal microscopy) and macroscopic characterization (swelling and rheology), we have confirmed that Olympic gelation takes place if and only if the plasmid library is enzymatically activated and sufficiently concentrated to allow random concatenation above the predicted ring overlap concentration. We observe a unique concentration-dependent trend in swelling that is expected for entangled gel networks but not for gels connected by conventional network junctions. Moreover, the shear modulus at frequencies well below the relaxation time of the rings develops a scaling that is anticipated for entanglement dominated systems and clearly distinct from conventional cross-linked gels that are dominated by the elastic contribution of network junctions. We anticipate that our current experimental data and future rheological studies into this new material type will contribute valuable information on the deformation behavior of purely entangled networks and help to improve existing models for rubber elasticity^50^.

Besides its function as a model system for ideal entangled networks, the creation of Olympic gels composed of genetic material directly proposes interesting potential applications for the fields of synthetic biology and bioengineering. In particular, sequences encoding for therapeutically relevant molecules, such as mRNA (e.g., for COVID or cancer vaccines), aptamers, or peptides can be included on the plasmid backbone. Owing to the free movement of the plasmid DNA, these molecules can be perpetually transcribed within Olympic gel particles via rolling circle amplification, in essence creating artificial microreactors^63^ that can deliver molecules at high concentration to specific target sites. Olympic gel particles could also act as artificial nuclei in synthetic cells in a similar manner as the topologically interlocked kinetoplast functions as the store of mitochondrial genes in *Kinetoplastea.* In the future, we will also explore Olympic gels as biotechnologically produced, sustainable, permeable, robust and yet biodegradable performance materials suitable for demanding applications such as ultrafiltration.

## Methods

### Initial thermodynamic prediction for the lock & key domain hybridization

To estimate thermodynamic parameters of the plasmid ring closure, an initial nearest-neighbor thermodynamic prediction was carried out for the 20 nt sticky end using the NUPACK software package (version 4.0.1.11)^41^. The prediction was based on the sequence GGN_16_CC, where N_16_ was represented by a randomized sequence (ACTTAGTCACTGTGGC) along with its reverse complement. For this pair of strands, the MFE values were calculated using the equation MFE_calculated_ = MFE_x,y_ – MFE_x_ – MFE_y_, where MFE_x,y_ is the predicted minimum free energy of the hybridized product, and MFE_x_ and MFE_y_ are the secondary structure energies of each of the two strands, respectively. The model parameters were set to T = 20 °C and 50 mM NaCl, with a maximum complex size set to 2. MFE was predicted as −114 kJ mol^−1^ (∼46 k_B_T).

### PVS10-DLK plasmid cloning

Cloning of the PVS10-DLK plasmid library was performed using a PVS10 plasmid (AddGene #104398) and a synthetically-produced single-stranded DNA oligonucleotide containing mixed “N” bases (termed the “DLK insert”, acttgtcagcgactgaggaaacagacNNNNNNNNNGCTGAGGNNNNNNNNNNNNNNNNCCTCAGCNNNNNNNNNtgcaggaaatgctcaccgttaagtct, synthesized by Integrated DNA Technologies) as follows. The PVS10 plasmid was digested with NcoI-HF and SbfI-HF enzymes according to recommended protocol (New England Biolabs). The resulting linear fragment was purified from an agarose gel using a gel extraction kit (Qiagen). The DLK insert oligonucleotide was inserted into the digested and purified PVS10 fragment via Gibson assembly using the NEBuilder® HiFi DNA Assembly Master Mix (New England Biolabs) to create the PVS10-DLK plasmid. The sequences of the PVS10 plasmid and PVS10-DLK plasmid are included in Supplementary Data 1, arbitrarily beginning at the start of the first DLK homology arm.

### Large-scale plasmid preparation

The Gibson-assembled PVS10-DLK plasmid library (2 μL) was transformed into at least five separate 50 μL aliquots of 5-alpha *E. coli* cells (New England Biolabs). Note that increasing the number of separate transformations from 5 to 20 increased the number of bacterial cells and therefore resulted in higher sequence diversity (Supplementary Figure S11). Each transformation was inoculated into 3 mL of LB media supplemented with 100 µg mL^−1^ ampicillin, grown for 8 hours at 37 °C with shaking at 200 rpm, and then mixed together and inoculated into 0.45 L of LB media per transformation supplemented with 100 µg mL^−1^ ampicillin. The culture was outgrown for 16–18 hours at 37 °C with shaking at 200 rpm. The bacterial culture was then centrifuged at 10,000 xg for 10 minutes. After discarding the supernatant, plasmid DNA was extracted and purified from bacterial pellets over a ZymoPURE™ II Plasmid Gigaprep column (Zymo Research) and recovered in nuclease-free water.

### Olympic gel preparation

A portion of the purified PVS10-DLK solution was nicked with Nt.BbvCI enzyme (New England Biolabs) for 16 hours at 37 °C to create the “Olympic” plasmid sample. A second portion was nicked with Nb.BsmI enzyme (New England Biolabs) for 4 hours at 65°C to create the relaxed “control” plasmid sample. To confirm successful nicking, a digestion test was performed on 1 μg of Nt.BbvCI-nicked and Nb.BsmI-nicked PVS10-DLK with HindIII-HF enzyme (New England Biolabs). Digestion products were run on a 1% w/v alkaline denaturing agarose gel^64^ to confirm the correct digestion pattern. We confirmed the successful nicking of the Olympic sample, which shows two bands indicative of the linearization of the PVS10 backbone via a double-strand break by HindIII, followed by dissociation of the nicked lock & key domain in the DLK insert. Successful nicking of the control sample was expected to result in three bands, a full-length single-stranded plasmid and two shorter single-stranded nicked fragments (Figure 2a). The control samples contained a majority of relaxed, circular plasmids (91%), with a small portion of supercoiled plasmids remaining (9%) (Supplementary Figure S6) . After confirmation of successful nicking, Nt.BbvCI, Nb.BsmI and recombinant Albumin (present in the enzymatic buffers) were degraded through enzymatic digestion with Proteinase K (New England Biolabs) for 16 hours at 37 °C. PVS10-DLK samples were then dialyzed through a 100 kDa cutoff membrane filter to remove protein and adjust buffer conditions to 2 mM Tris-HCl, 0.1 mM EDTA, 2.5 mM NaCl, pH 8.0. Samples were then further concentrated according to the desired test concentrations by reducing the volume as required by vacuum evaporation at room temperature. Buffer conditions for all samples were adjusted to a final composition of 20 mM Tris-HCl, 2 mM EDTA, 50 mM NaCl, pH 8.0 (referred to as Tris-NaCl buffer). To create the Olympic gel, Nt.BbvCI-nicked PVS10-DLK solutions were incubated at 50 °C for 30 minutes to transiently open the plasmids, then slowly cooled by −3 °C per minute to 20 °C to anneal plasmids into an interlinked gel. Control samples were treated in the same manner.

### Gel-shift assay

In order to confirm proper concatenation, a gel-shift assay was performed on control and Olympic samples prepared and annealed at 0.01 wt%, 0.025 wt%, 0.05 wt%, 0.075 wt%, 0.1 wt%, 0.125 wt%, and 0.25 wt%. Shortly before sample loading into the gel, annealed samples were diluted in Tris-NaCl buffer to 5 ng μL^−1^. 50 ng of each sample were loaded in a 0.5% w/v native agarose gel, run at 100 V for 2 hours, and imaged on a Typhoon™ FLA 9500 imager.

### Next-generation sequencing

The Gibson assembly product produced before transformation into *E. coli* and the final PVS10-DLK plasmid library isolated after large-scale amplification and purification were sequenced using NGSelect Amplicon 2nd PCR sequencing (Eurofins Genomics) on the Illumina MiSeq. The sequences were merged using FLASH (v 2.2.00)^65^ and subsequently converted to FASTA format. A custom python script was used to identify instances of Nt.BbvCI nicking sites (GCTGAGG & CCTCAGC), extract sequences of interest, and identify unique sequence variants and their frequencies (Supplementary Data 2). The base-wise variation was analyzed by generating a base-wise frequency matrix and plotted using Logomaker^66^. Increasing the number of parallel transformations from 5 to 20 increased the number of distinct sequence variants in the final library from 16,307 to 26,118 (Supplementary Figure S11b–d). However, this increase in sequence complexity is not required from a theoretical point of view.

### Thermodynamic simulations based on sequencing data

The full library of 16,307 DLK sequences (lock & key domains as GGN_16_CC) generated from next-generation sequencing were arranged in descending order of prevalence. A library containing reverse complements of these sequences was generated using Biopython in a custom python script (Supplementary Data 3). The MFE values for all explicit lock & key pairs were calculated for all possible lock & key combinations in the library with the NUPACK software package (version 4.0.1.11)^41^. For each lock & key pair, the MFE values were calculated using the equation MFE_calculated_ = MFE_x,y_ – MFE_x_ – MFE_y_, where MFE_x,y_ is the predicted minimum free energy of the hybridized product, and MFE_x_ and MFE_y_ are the secondary structure energies of each of the two strands, respectively. The computations were performed on the Barnard CPU cluster at the Zentrum für Informationsdienste und Hochleistungsrechnen (ZIH), Technische Universität Dresden. Post computation, the MFE values were merged and processed using a custom python script. The relative frequency, p(x), of each unique variant, x, was calculated as p(x) = n/N, where n represents the frequency of each unique variant, and N denotes the total number of sequences obtained from the next-generation sequencing data. The Simpson’s Diversity Index (S)^67^ was calculated using the equation: S= Σp(x)². The Shannon Information Entropy (H)^68^ was calculated using the equation: H=-Σ p(x) × log_2_(p(x)).

### Atomic Force Microscopy

Sample preparation protocols were adapted from Refs. [^69, 70^]. Control and Olympic samples were prepared and annealed at a concentration below *c_gel_* (0.03–0.075 wt%). After annealing, the samples were stored at 20 °C overnight. The samples were diluted to 5 ng μL^−1^ (0.0005 wt%) in Tris-NaCl buffer immediately prior to imaging. The dilution was done to ensure that multiple plasmids were unlikely to adsorb to the same spot on the mica surface unless they were physically linked together. A freshly cleaved mica disc was covered with a 25 μL droplet of 10 mM Tris-HCl, 25 mM MgCl_2_, pH 8.0, then 5 ng of the PVS10-DLK sample was pipetted into the droplet and mixed with further pipetting. The disc was incubated for 5 minutes at room temperature in an enclosed container with a humid environment to reduce evaporation. After incubation, the sample was washed three times by addition of 50 μL of 10 mM Tris-HCl, 3 mM NiCl_2_, pH 8.0 (Tris-NiCl_2_ buffer) followed by vigorous pipetting and removal of the same volume. An additional 50 μL of Tris-NiCl_2_ buffer was added to the sample prior to imaging. AFM imaging was performed in Tris-NiCl_2_ buffer. Peak Force Tapping imaging was performed on a Dimension FastScan AFM system (Bruker) in the ScanAsyst in Fluid - HiRes mode using Fastscan-D-SS high-resolution cantilevers (Bruker). Peak Force amplitude was set to 10 nm at a frequency of 8000 Hz. The Peak Force setpoint was set to 0.02 V. High resolution images of adsorbed plasmids were taken at a scan rate of 2.82 Hz and 1024 samples/line.

### Image analysis on Atomic Force Microscopy data

AFM images were analyzed in Fiji^71^ (version 1.54m). Images were converted to 8-bit, then a Gaussian blur filter with a sigma of 2.0 was applied. A threshold was applied to mask objects in the image over a dark background. To remove background noise, particles with a circularity above 0.2–0.5 (depending on background noise levels) were removed through masking via the “Analyze Particles” menu. The mask was converted to binary and skeletonized. The total branch length for each skeletonized object (i.e. the total contour length of the DNA) was measured using the “Particle Analyser” feature from the BoneJ plugin^72^ (version 1.4.3).

### Cryo-Scanning Electron Microscopy

Cryo-SEM imaging was performed on a Zeiss Crossbeam 550 SEM fitted with a Quorum nitrogen gas-cooled cold stage for imaging of samples under cryogenic conditions. Olympic gel, control, and non-nicked samples were prepared and annealed at 0.25 wt%. 2 μL of sample were subjected to high-pressure freezing^49^ on a Leica EM ICE High Pressure Freezer and subsequently stored in liquid nitrogen. A Quorum PP3010T cryogenic preparation system was used to mount samples under liquid nitrogen and vacuum conditions for subsequent transfer to the cryo-preparation chamber and loading onto the pre-cooled SEM stage. Sample surfaces were imaged under vacuum without etching or coating at low acceleration voltage and probe current (1.8 kV and 25 pA respectively) at 13.8k x magnification. Samples were sublimed by slowly warming the SEM stage to −90 °C, holding at −90 °C for 10 minutes, then cooling to −145 °C for imaging.

### Computer simulations

Ideal Olympic gels made of mono-disperse flexible polymer rings were prepared as described in Ref. [^19^] using the bond fluctuation model (BFM)^73,74^ in the LeMonADE implementation on graphical processing units (GPU)^75,76^. A polymer volume fraction of *φ* = 0.5 was chosen on a cubic lattice of size 256^3^. The degrees of polymerization of the cyclic polymers were *N* = 64, 128, 256, 512, and 1024. Shear simulations were performed as described in Refs. [^53,77^]. Details of the simulations and the analysis are discussed in Supplementary Section S1.

### Tube inversion test

Olympic gel, control, and non-nicked samples (100 µL each) were prepared at a concentration of 0.25 wt% in DNA-low-bind Eppendorf tubes. The plasmid samples were first centrifuged with a desktop microcentrifuge. Subsequently, the tubes were inverted upside down for 1 minute and a photo was taken. The content of the tubes was then centrifuged again and the tubes were annealed at 50 °C for 30 minutes, then slowly cooled to 20 °C at a rate of −3 °C min^−1^. After annealing, the tubes were inverted again and a second photo was taken after 1 minute.

### Swelling measurements

Control and Olympic samples were prepared and annealed at 0.025 wt%, 0.05 wt%, 0.125 wt%, 0.25 wt%, 0.5 wt%, and 0.8 wt% in the presence of 200x diluted SYBR Gold (Thermo Fisher, cat. #S11494) prior to the measurement. Shortly before imaging, 5 μL of each sample were dispensed into a glass-bottom 384-well plate (Greiner Bio-One). Each sample was immersed in 100 μL of Tris-NaCl buffer. Images were acquired every hour for 48 hours at 4x magnification on the Andor Dragonfly confocal microscope (Oxford Instruments). The samples were kept in buffer for one week and then centrifuged at 300 xg for 2 minutes before the end-point images were taken. The gel volume was quantified using IMARIS software (Oxford Instruments) via the surface wizard under a uniform threshold.

### Rheology

Viscoelasticity measurements were conducted using an Anton Paar MCR301 rheometer with a 25 mm diameter cone-plate geometry (cone angle 0.5°). Temperature control was ensured by installing a Peltier system PTD-200 (Anton Paar) within the measurement area. The chamber is equipped with a solvent trap and a vapor lock to prevent evaporation artifacts. A wet paper cylinder was placed around the plate’s circumference to maintain high humidity inside the chamber. The annealing protocol started with holding the PVS10-DLK solution at equilibrium at 20 °C for 5 minutes, followed by heating steps from 20 °C to 50 °C at 3 °C min^−1^, and then incubation at 50 °C for 30 minutes. The samples were then cooled to 20 °C at −3 °C min^−1^ and held at 20 °C for 120 minutes, all at 1% strain and 1 Hz. Frequency sweeps were conducted from 0.001 Hz to 10 Hz at 1% strain for 2 measurement repeats, while amplitude sweeps were performed from 0.1% to 1000% strain at 1 Hz for 4 measurement repeats. Frequency sweeps for the lowest concentrations approached the instrument’s sensitivity limit. Collected data with torque values below the threshold of 5 nNm was therefore excluded from the plots. Stress-relaxation properties were measured at 15% strain, where the strain was kept constant while recording shear stress over time. Stress-strain curves were determined by shearing rotationally at increased shear rate from 0.0001 s^−1^ to 1 s^−1^ as the respective shear stress was recorded.

## Acknowledgements

E.K. acknowledges funding by the Federal Ministry of Education and Research of Germany (BMBF) in the program NanoMatFutur (grant no. 13XP5098). M.L. acknowledges funding by the Deutsche Forschungsgemeinschaft (DFG, grant #329888557). J.-U.S. was supported by the DFG under Germany’s Excellence Strategy (EXC2068−390729961, Cluster of Excellence Physics of Life of TU Dresden). The authors acknowledge the computing time and support on the high-performance computer at the Nationales Hochleistungs Rechnen (NHR) Center of TU Dresden. The authors thank Dr. Lothar Steeb and Pelobiotech GmbH for supporting us as an industry partner. The authors thank Prof. Dr. Carsten Werner for useful discussions, Qiong Li and Jannes Dressel for assistance with AFM measurements, Vinolia Dmello for cryo-SEM sample preparation, Dr. Luca Bertinetti and Prof. Dr. Yael Politi for assistance with cryo-SEM measurements and data analysis, Dr. Günter Auernhammer for helpful discussions on rheology experiments, Srividhya Sainath and Apurv Deepak Kulkarni for initial support in setting up and scaling computational simulations, and Dr. Lars Renner, Dr. Stephanie Klinghammer, and Prof. Gianaurelio Cuniberti for providing access to microbiology facilities. We also thank Dr. Michele Marass for giving valuable feedback on the draft. Parts of Figure 1d were created with BioRender.com.

## Author contributions

E.K., J.-U.S. and M.L. conceived and designed the study. A.A. and S.K.S. designed, carried out, and analyzed cloning experiments and production of plasmids for the Olympic gel and control samples. S.K.S. and I.H. performed the AFM measurements. Y.-H.P. performed the rheological experiments, swelling measurements, and the gel-shift assay. K.G. carried out production of plasmids for the Olympic gel and control samples, next-generation sequencing experiments and data analysis, thermodynamic calculations, and participated in the design of cloning experiments. M.L. and T.M. carried out computer simulations. C.F. performed the cryo-SEM experiments. S.K.S., A.A., Y.-.H.P., K.G., M.L., J.-U.S. and E.K. participated in data analysis. M.L. wrote the supporting information with input from all authors. S.K.S., Y.-H.P., M.L., and E.K. wrote the manuscript with input from all authors.

## Data availability

All data supporting the findings of this study are provided in the article and its Supplementary Information.

## Code availability

Code used in the analysis of next-generation sequencing data is provided as Python scripts in the Supplementary Information. Software that can be used to reproduce the simulation results is available at Zenodo (https://doi.org/10.5281/zenodo.5513487 and https://doi.org/10.5281/zenodo.5061542).

## References

1. Osada, Y. & Gong, J.-P. Soft and Wet Materials: Polymer Gels. Adv. Mater. 10, 827– 837 (1998).

2. Sangeetha, N. M. & Maitra, U. Supramolecular gels: Functions and uses. Chem. Soc. Rev. 34, 821–836 (2005).

3. Hirst, A. R., Escuder, B., Miravet, J. F. & Smith, D. K. High-Tech Applications of Self-Assembling Supramolecular Nanostructured Gel-Phase Materials: From Regenerative Medicine to Electronic Devices. Angew. Chem. Int. Ed. 47, 8002–8018 (2008).

4. Qiu, Y. & Park, K. Environment-sensitive hydrogels for drug delivery. Adv. Drug Deliv. Rev. 53, 321–339 (2001).

5. Kim, J. et al. Wearable smart sensor systems integrated on soft contact lenses for wireless ocular diagnostics. Nat. Commun. 8, 14997 (2017).

6. Lee, K. Y. & Mooney, D. J. Hydrogels for Tissue Engineering. Chem. Rev. 101, 1869– 1880 (2001).

7. Krieg, E., Weissman, H., Shirman, E., Shimoni, E. & Rybtchinski, B. A recyclable supramolecular membrane for size-selective separation of nanoparticles. Nat. Nanotechnol. 6, 141–146 (2011).

8. Beebe, D. J. et al. Functional hydrogel structures for autonomous flow control inside microfluidic channels. Nature 404, 588–590 (2000).

9. Krieg, E. et al. Supramolecular Gel Based on a Perylene Diimide Dye: Multiple Stimuli Responsiveness, Robustness, and Photofunction. J. Am. Chem. Soc. 131, 14365–14373 (2009).

10. Flory, P. J. Principles of Polymer Chemistry. (Cornell University Press, Ithaca, NY, 1953).

11. 11. De Gennes, P. G. Scaling Concepts in Polymer Physics. (Cornell University Press, Ithaka and London, 1979).

12. Hart, L. F. et al. Material properties and applications of mechanically interlocked polymers. Nat. Rev. Mater. 6, 508–530 (2021).

13. Vilgis, T. A. & Otto, M. Elasticity of entangled polymer loops: Olympic gels. *Phys*. Rev. E 56, R1314–R1317 (1997).

14. Lang, M., Fischer, J., Werner, M. & Sommer, J.-U. Swelling of Olympic Gels. Phys. Rev. Lett. 112, 238001 (2014).

15. Shapiro, T. A. & Englund, P. T. The Structure and Replication of Kinetoplast Dna. Annu. Rev. Microbiol. 49, 117–143 (1995).

16. Chen, J., Rauch, C. A., White, J. H., Englund, P. T. & Cozzarelli, N. R. The topology of the kinetoplast DNA network. Cell 80, 61–69 (1995).

17. He, P., Katan, A. J., Tubiana, L., Dekker, C. & Michieletto, D. Single-Molecule Structure and Topology of Kinetoplast DNA Networks. Phys. Rev. X 13, 021010 (2023).

18. Hoffmann, A., et al. Molecular model of the mitochondrial genome segregation machinery in Trypanosoma brucei. Proc. Natl. Acad. Sci. 115, E1809–E1818 (2018).

19. Fischer, J., Lang, M. & Sommer, J.-U. The formation and structure of Olympic gels. J. Chem. Phys. 143, 243114 (2015).

20. Ubertini, M. A. & Rosa, A. Computer simulations of melts of ring polymers with nonconserved topology: A dynamic Monte Carlo lattice model. *Phys*. Rev. E 104, 054503 (2021).

21. Bonato, A., Marenduzzo, D., Michieletto, D. & Orlandini, E. Topological gelation of reconnecting polymers. Proc. Natl. Acad. Sci. 119, e2207728119 (2022).

22. Stoddart, J. F. Mechanically Interlocked Molecules (MIMs)—Molecular Shuttles, Switches, and Machines (Nobel Lecture). Angew. Chem. Int. Ed. 56, 11094–11125 (2017).

23. Lu, C.-H., et al. Switchable Reconfiguration of an Interlocked DNA Olympiadane Nanostructure. Angew. Chem. Int. Ed. 53, 7499–7503 (2014).

24. Datta, S. et al. Self-assembled poly-catenanes from supramolecular toroidal building blocks. Nature 583, 400–405 (2020).

25. Bardot, M. I. et al. Mechanically interlocked two-dimensional polymers. Science 387, 264–269 (2025).

26. Lang, M. & Kumar, K. S. Simple and General Approach for Reversible Condensation Polymerization with Cyclization. Macromolecules 54, 7021–7035 (2021).

27. Pickett, G. T. DNA-origami technique for olympic gels. Europhys. Lett. EPL 76, 616– 622 (2006).

28. Raphaifl, E., Gay, C. & De Gennes, P. G. Progressive Construction of an ‘Olympic’ Gel. J. Stat. Phys. 89, 111–118 (1997).

29. Endo, K. & Yamanaka, T. Copolymerization of Lipoic Acid with 1,2-Dithiane and Characterization of the Copolymer as an Interlocked Cyclic Polymer. Macromolecules 39, 4038–4043 (2006).

30. Endo, K., Shiroi, T., Murata, N., Kojima, G. & Yamanaka, T. Synthesis and Characterization of Poly(1,2-dithiane). Macromolecules 37, 3143–3150 (2004).

31. Xiong, X. et al. Evaporation-Assisted Synthesis of Olympic Gels. Angew. Chem. Int. Ed. e202425034 (2025) doi:10.1002/anie.202425034.

32. Krajina, B. A., Zhu, A., Heilshorn, S. C. & Spakowitz, A. J. Active DNA Olympic Hydrogels Driven by Topoisomerase Activity. Phys. Rev. Lett. 121, 148001 (2018).

33. Kim, Y. S. et al. Gelation of the genome by topoisomerase II targeting anticancer agents. Soft Matter 9, 1656–1663 (2013).

34. Note: Ref. 32 demonstrates that Topoisomerase II causes a notable degree of plasmid concatenations. However, their data also shows that (1) a major fraction of plasmids likely remained non-concatenated (Suppl. Fig. S2 in ref. 32), and (2) the complex modulus (G*) drastically decreases when the material is exposed to Proteinase K, an enzyme that digests Topoisomerase II (Suppl. Fig. S3 in ref. 32). The latter observation highlights that Topoisomerase II itself forms classical crosslinks that dominate the gel’s mechanical characteristics, as the authors correctly note. The authors assume that the digestion of Topoisomerase II by Proteinase K is complete and the remaining G* is therefore a result of a purely topological network. However, the completeness of the digestion was not demonstrated, and appears unlikely, since the presence of EDTA in the reaction (Suppl. Sect. I.F in ref. 32) would have a strong inhibitory effect on Proteinase K.

35. Peng, Y.-H. et al. Dynamic matrices with DNA-encoded viscoelasticity for cell and organoid culture. Nat. Nanotechnol. 18, 1463–1473 (2023).

36. Speed, S. K., Gupta, K., Peng, Y.-H., Hsiao, S. K. & Krieg, E. Programmable polymer materials empowered by DNA nanotechnology. J. Polym. Sci. 61, 1713–1729 (2023).

37. Wong, C.-H. & Zimmerman, S. C. Orthogonality in organic, polymer, and supramolecular chemistry: from Merrifield to click chemistry. Chem. Commun. 49, 1679– 1695 (2013).

38. Krieg, E., Bastings, M. M. C., Besenius, P. & Rybtchinski, B. Supramolecular Polymers in Aqueous Media. Chem. Rev. 116, 2414–2477 (2016).

39. Xu, Q., Schlabach, M. R., Hannon, G. J. & Elledge, S. J. Design of 240,000 orthogonal 25mer DNA barcode probes. Proc. Natl. Acad. Sci. 106, 2289–2294 (2009).

40. Gibson, D. G. et al. Enzymatic assembly of DNA molecules up to several hundred kilobases. Nat. Methods 6, 343–345 (2009).

41. Fornace, M. E. et al. NUPACK: Analysis and Design of Nucleic Acid Structures, Devices, and Systems. Preprint at 10.26434/chemrxiv-2022-xv98l (2022).

42. Lang, M., Fischer, J. & Sommer, J.-U. Effect of Topology on the Conformations of Ring Polymers. Macromolecules 45, 7642–7648 (2012).

43. Diao, Y., Rodriguez, V., Klingbeil, M. & Arsuaga, J. Orientation of DNA Minicircles Balances Density and Topological Complexity in Kinetoplast DNA. PLOS ONE 10, e0130998 (2015).

44. Svetlov, V. & Artsimovitch, I. Purification of Bacterial RNA Polymerase: Tools and Protocols. in Bacterial Transcriptional Control: Methods and Protocols (eds. Artsimovitch, I. & Santangelo, T. J.) 13–29 (Springer, New York, NY, 2015). doi:10.1007/978-1-4939-2392-2_2.

45. Harnett, J., Weir, S. & Michieletto, D. Effects of monovalent and divalent cations on the rheology of entangled DNA. Soft Matter 20, 3980–3986 (2024).

46. Levy, M. S. et al. Effect of shear on plasmid DNA in solution. Bioprocess Eng. 20, 7–13 (1999).

47. Integrated DNA Technologies. Mixed bases | IDT. Integrated DNA Technologies https://eu.idtdna.com/pages/products/custom-dna-rna/mixed-bases.

48. Menger, F. M., Seredyuk, V. A., Apkarian, R. P. & Wright, E. R. Colloidal Assemblies of Branched Geminis Studied by Cryo-etch-HRSEM. J. Am. Chem. Soc. 124, 12408–12409 (2002).

49. Aston, R., Sewell, K., Klein, T., Lawrie, G. & Grøndahl, L. Evaluation of the impact of freezing preparation techniques on the characterisation of alginate hydrogels by cryo-SEM. Eur. Polym. J. 82, 1–15 (2016).

50. Rubinstein, M. & Panyukov, S. Elasticity of Polymer Networks. Macromolecules 35, 6670–6686 (2002).

51. Ball, R. C., Doi, M., Edwards, S. F. & Warner, M. Elasticity of entangled networks. Polymer 22, 1010–1018 (1981).

52. Edwards, S. F. & Vilgis, Th. The effect of entanglements in rubber elasticity. Polymer 27, 483–492 (1986).

53. Rubinstein, M. & Colby, R. Polymer Physics. (Oxford University Press, Oxford, United Kingdom, 2004).

54. Yamamoto, T., Campbell, J. A., Panyukov, S. & Rubinstein, M. Scaling Theory of Swelling and Deswelling of Polymer Networks. Macromolecules 55, 3588–3601 (2022).

55. Kapnistos, M. et al. Unexpected power-law stress relaxation of entangled ring polymers. Nat. Mater. 7, 997–1002 (2008).

56. Doi, Y. et al. Melt Rheology of Ring Polystyrenes with Ultrahigh Purity. Macromolecules 48, 3140–3147 (2015).

57. Regan, K., Ricketts, S. & Robertson-Anderson, R. M. DNA as a Model for Probing Polymer Entanglements: Circular Polymers and Non-Classical Dynamics. Polymers 8, 336 (2016).

58. Paiva, W. A. et al. From Bioreactor to Bulk Rheology: Achieving Scalable Production of Highly Concentrated Circular DNA. Adv. Mater. 36, 2405490 (2024).

59. Peddireddy, K. R. et al. Unexpected entanglement dynamics in semidilute blends of supercoiled and ring DNA. Soft Matter 16, 152–161 (2020).

60. Michieletto, D. et al. Topological digestion drives time-varying rheology of entangled DNA fluids. Nat. Commun. 13, 4389 (2022).

61. Miracle, D. B. & Senkov, O. N. A critical review of high entropy alloys and related concepts. Acta Mater. 122, 448–511 (2017).

62. Yeh, J. W. Recent progress in high-entropy alloys. Ann. Chim. Sci. Mater. Paris 31, 633–648 (2006).

63. Hahn, J., Chou, L. Y. T., Sørensen, R. S., Guerra, R. M. & Shih, W. M. Extrusion of RNA from a DNA-Origami-Based Nanofactory. ACS Nano 14, 1550–1559 (2020).

64. Green, M. R. & Sambrook, J. Alkaline Agarose Gel Electrophoresis. Cold Spring Harb. Protoc. 2021, pdb.prot100438 (2021).

65. Magoč, T. & Salzberg, S. L. FLASH: fast length adjustment of short reads to improve genome assemblies. Bioinforma. Oxf. Engl. 27, 2957–2963 (2011).

66. Tareen, A. & Kinney, J. B. Logomaker: beautiful sequence logos in Python. Bioinformatics 36, 2272–2274 (2020).

67. Simpson, E. H. Measurement of Diversity. Nature 163, 688–688 (1949).

68. Shannon, C. E. A mathematical theory of communication. Bell Syst. Tech. J. 27, 379– 423 (1948).

69. Pyne, A., Thompson, R., Leung, C., Roy, D. & Hoogenboom, B. W. Single-Molecule Reconstruction of Oligonucleotide Secondary Structure by Atomic Force Microscopy. Small 10, 3257–3261 (2014).

70. Holmes, E. P. et al. Under or Over? Tracing Complex DNA Structures with High Resolution Atomic Force Microscopy. Preprint at 10.1101/2024.06.28.601212 (2024).

71. Schindelin, J., et al. Fiji: an open-source platform for biological-image analysis. Nat. Methods 9, 676–682 (2012).

72. Doube, M. et al. BoneJ: Free and extensible bone image analysis in ImageJ. Bone 47, 1076–1079 (2010).

73. Carmesin, I. & Kremer, K. The bond fluctuation method: a new effective algorithm for the dynamics of polymers in all spatial dimensions. Macromolecules 21, 2819–2823 (1988).

74. Deutsch, H. P. & Binder, K. Interdiffusion and self-diffusion in polymer mixtures: A Monte Carlo study. J. Chem. Phys. 94, 2294–2304 (1991).

75. Müller, T., Wengenmayr, M., Dockhorn, R. & Knespel, M. LeMonADE-project/LeMonADE-GPU: Release v1.2. Zenodo 10.5281/zenodo.5513487 (2021).

76. Müller, T., et al. LeMonADE-project/LeMonADE: LeMonADE v2.2.2. Zenodo 10.5281/zenodo.5061542 (2021).

77. Lang, M. & Müller, T. On the Reference Size of Chains in a Network and the Shear Modulus of Unentangled Networks Made of Real Chains. Macromolecules 55, 8950–8959 (2022).

