## Supplementary Information for "Assembling a true ‘Olympic Gel’ from >16,000 combinatorial DNA rings"

### S1. COMPUTER SIMULATIONS

We apply the bond fluctuation model (BFM) [1, 2] to simulate Olympic gels made of flexible chains. All simulations discussed rely on the LeMonADE implementation [3, 4] of the BFM model on graphical processing units (GPU). Olympic gels were prepared at a polymer volume fraction of $\phi = 0.5$ on a cubic lattice with $L^3 = 256^3$ lattice sites and periodic boundary conditions leading to $2^{20}$ monomers per samples. In a monodisperse solution of freely concatenating ring polymers, the average number of concatenations per ring, $f_n$, is proportional to $\phi R_g^2$ [5], which provides $f_n \propto \phi N$ for rings with ideal conformations (see section S2 for a more detailed discussion). Here, $R_g$ is the radius of gyration of the rings and $N$ is the number of Kuhn segments of the ring. $f_n$ controls the formation of Olympic gels [6]. Thus, one can study gelation by varying $\phi$ or $N$ or both. The plateau modulus of entangled samples is largest for the highest (melt) concentrations providing the best signal to noise ratio in computer simulations, in particular, at small deformations, see Figure S1 for illustration. Therefore, concatenated ring polymer melts (perfectly monodisperse, no linear chains) were prepared with degrees of polymerization of $N = 64$, 128, 256, 512, 1024 as described in Ref. [6]. Shear simulations in $x$ direction within the $xz$ plane were performed as discussed in detail in Refs. [7, 8] by applying a small external force $\vec{f}$ to all monomers located on several selected layers of the sample, see also Figure 4a for illustration (All cross-referenes to Figures and sections starting with S refer to the supporting information, all other cross-references concern the main document).

From the beginning of the deformation, the trajectories of all particles are followed. This allows to compute the average displacement $\Delta x$ in $x$ direction of all monomers in a given $z$ layer. The latter provides the shear strain $\gamma = 2\Delta x/L$ averaged over the two halves of the sample, see Figure 3 of Ref. [8]. Shear strain is determined throughout the full simulation of the shear deformation process as shown in Figure S1 and

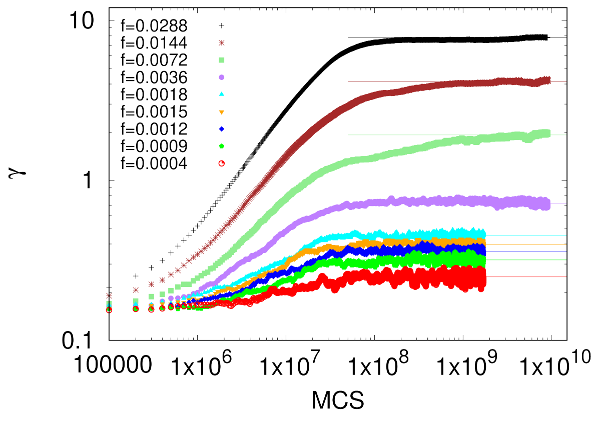


FIG. S1. Shear strain $\gamma$ as a function of simulation time given in Monte Carlo steps (MCS) for a series of applied shear forces, $f$, for networks with $N = 1024$.

the average shear strain is determined in the equilibrium shear region. In the simulations, the lattice unit is $a$, the energy unit is $kT$ such that the unit of force is $kT/a$ and stress is given in units of $kT/a^3$. The shear stress, $\sigma_{xy}$, is the total average force in $x$ direction, $f_x$, divided by the area $A_z = L^2$ of the $xy$ plane,

$$\sigma_{xz} = \frac{f_x}{A_z} \approx \frac{f}{8a^2} \quad \text{(S1)}$$

and plotted as a function of shear strain in Figure 4b and Figure S2. The last relation in the above equation reflects symmetry and the parameters of our simulations, see Ref. [8] for details. Typically, the modulus is determined in the linear deformation regime at small strains

$$G \approx \frac{\sigma_{xz}}{\gamma}. \quad \text{(S2)}$$

For the sample $N = 64$ close to the gelation threshold, the measured strain did not saturate as a function of time. Therefore, we cannot discuss the equilibrium elastic properties of this sample. For all samples $N \geq 128$, the measured strain saturates at large times, see Figures S1 and S3 for the most and the least concatenated samples as examples. Our simulation data provide a linear relationship $\sigma \propto \gamma$ at small strains only for the least concatenated sample $N = 128$, see Figure S2. The modulus of this sample falls clearly below the modulus of the more concatenated samples.

---

* Authors contributed equally.
†
‡

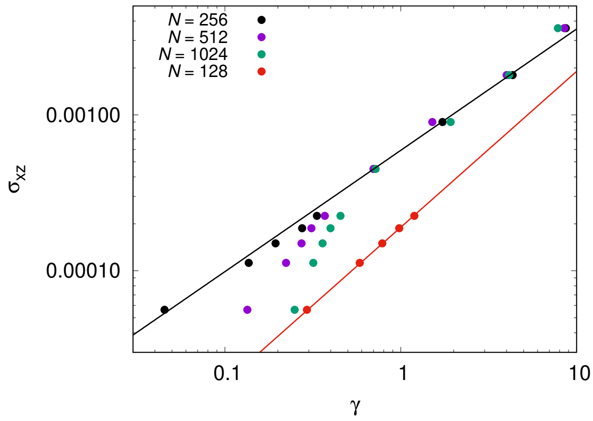


FIG. S2. Simulation results (dots) for the stress-strain relation of concatenated rings under shear stress $\sigma_{xz}$ for samples with varying $N$. The lines are fits by power laws (orange: $\sigma \propto \gamma$; black: $\sigma \propto \gamma^{0.78}$) as a function of strain $\gamma$.

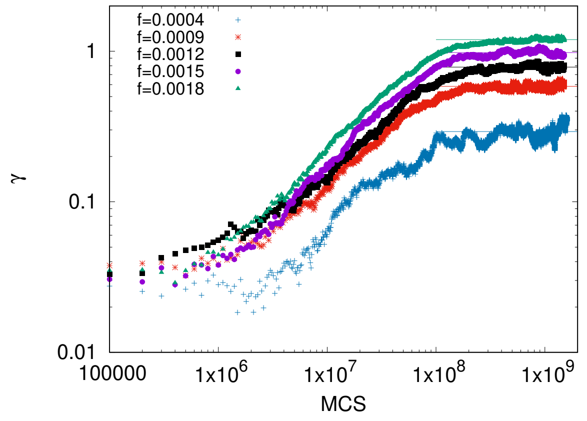


FIG. S3. $\gamma$ as a function of simulation time given in Monte Carlo steps (MCS) for a series of applied shear forces, $f$ for $N = 128$. The lines show the plateau level that was fit to all data at times beyond $5 \times 10^8$MCS.

The weight faction of the elastically active network strands is here around 70% (see the data for $\phi = 0.5$ at $f_w \approx 3.7$ in Figure 4 of Ref. [6]). Adopting standard models for entangled chains [9], we expect a reduction of modulus by a factor of roughly $0.7^{2.3} \approx 0.44$. However, the measured modulus at small $\gamma$ is clearly below this estimate. A possible explanation for this discrepancy stems from the network structure close to gelation. The majority of rings inside the elastically active material of sample $N = 128$ has exactly two elastically active concatenations. Thus, the elastic network structure forms predominantly poly[n]catenates between a small fraction of rings that serve as branching points. Poly[n]catenates are conformationally similar to ideal chains but more compact reducing the entanglement with the surrounding polymers [10]. Therefore, an additional reduction of modulus and a cross-over to a linear stress strain relation when approaching the gelation threshold is likely. Moreover, the sample contains about 10% of rings in the sol. These rings are subject to the applied force and cause a non-equilibrium contribution to the measured shear strain. A direct separation of these contributions is not possible as the orientation of the free rings couples to the orientation of the rings in the gel and the mobile rings exert a friction force onto the network. Therefore, we abstain from a quantitative analysis of the modulus for sample $N = 128$.

In Figure S2 (see also Figure 4b of the main text), we find for $N \geq 256$ universal relation of $\sigma \propto \gamma^{0.78 \pm 0.03}$ at $\gamma \gtrsim 1/2$ and below the dominance of finite extensibility starting around $\gamma \approx 5$. Here, the gels are nearly perfect containing virtually no rings that diffuse freely (see also Figure S4), and we expect no significant corrections due to network defects. In Figure 4b, the data at largest strains were excluded from the fit due to finite extensibility corrections and since the deformation starts to become non-homogeneous. The onset of a non-homogeneous deformation behavior is visible already for the longest chains in Figure S4, which was also excluded from the fit in Figure 4b (we checked all simulations visually to assure that $\gamma$ was averaged only over time intervals where the samples where deformed homogeneously).

The striking new observation is the super-linear dependence of $\sigma$ on $\gamma$ for the smallest applied forces in the limit of large $N$. Such a super-linear dependence was predicted in several models of entanglements for uni-axial deformation at low strains [11], however, the precise relation for shear was not discussed previously. This result shows that the rings may slip against each other in the limit of small strains. Remarkably, the effect of the slippage becomes more pronounced for larger $N$ that provide a longer contour length along which pairwise entanglements can slide. Such a qualitative tendency was discussed in slip link models for increasing $N$ between network junctions [12, 13]. Beyond a strain of $\gamma \approx 0.5$, the different slip as a function of $N$ becomes irrelevant as the deformation of the tubes starts to dominate leading to a universal deformation behavior below the onset of finite extensibility effects. For such intermediate strains, a sublinear stress-strain relation is expected for entangled chains and was discussed in detail only for the case of uni-axial extension [11, 14]. The above scaling relation refers to a weaker effective strain softening as proposed theoretically, which might be caused by a continuously increasing finite extensibility contribution. The data collapse for a weight average number of concatenations, $f_w$, of roughly $f_w \gtrsim 6$ when considering the data of Ref. [6]. A similar collapse of stress-strain data beyond a threshold overlap of the cyclic molecules could provide, therefore, a rough estimate of $f_w$ for a series of Olympic gels.

### S2. CONCENTRATION THRESHOLD FOR THE FORMATION OF OLYMPIC GELS

Preceding simulation studies [5, 6] have established that Olympic gels are only formed beyond a certain overlap of the concatenating cyclic molecules at preparation conditions. The corresponding threshold must be estimated for planning and analyzing the experi-

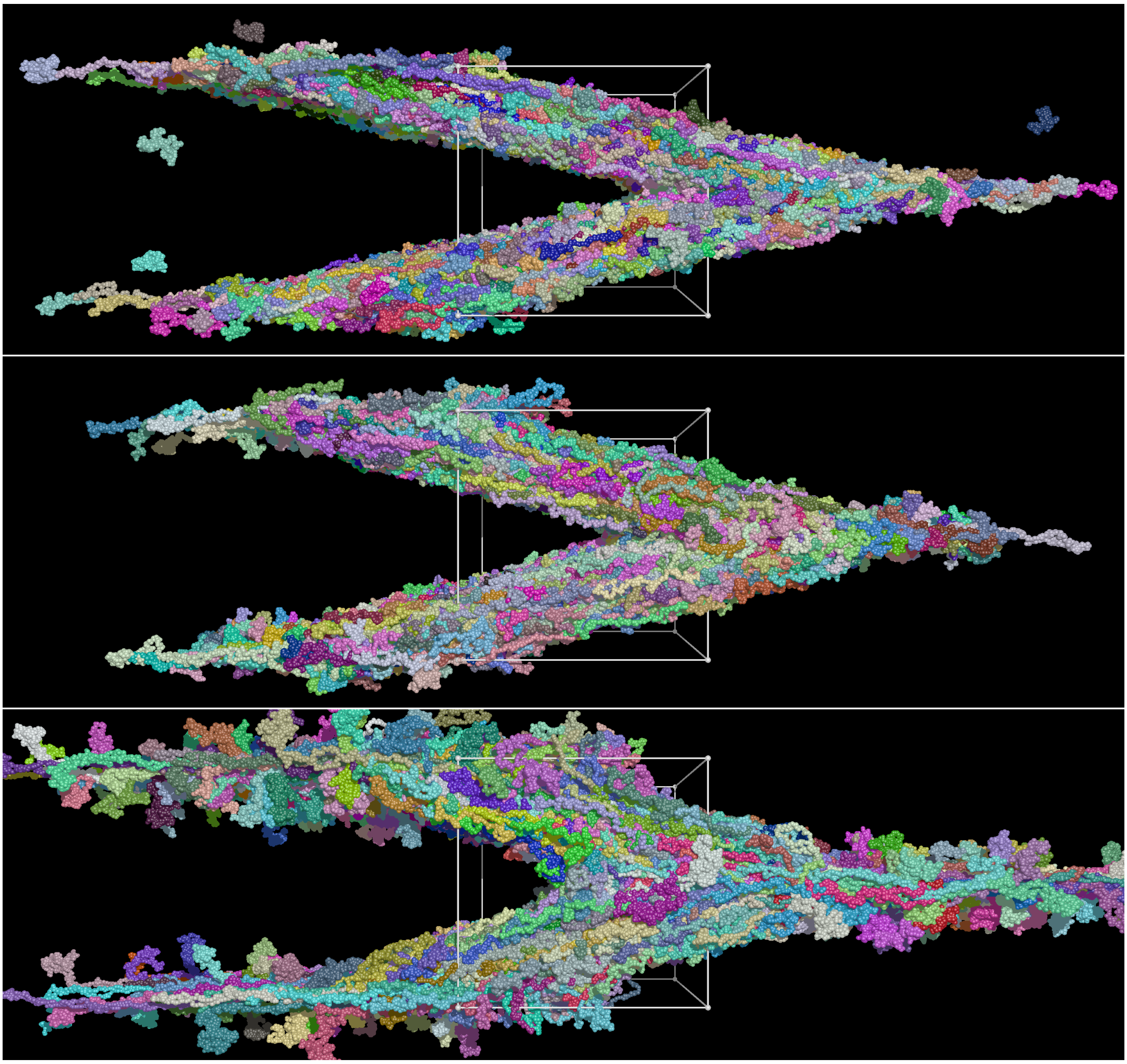

FIG. S4. Snapshots of $N = 256$, $N = 512$, $N = 1024$ at the second largest applied force. The deformation starts to becomes non-homogeneous for the largest $N$ at this force.

ments.

In Figure S5, the simulation data of Ref. [5] was replotted as a function of the overlap number

$$P = \frac{\phi R_g^3}{N v_0} \tag{S3}$$

that counts the average number of cyclic polymers in the pervaded volume of a polymer. Here, $R_g$ is the radius of gyration of the cyclic polymers, $\phi$ is the polymer volume fraction, $N$ is the number of Kuhn segments per polymer, and $v_0$ is the volume per Kuhn segment. As basis for this plot, only the data at a polymer volume fraction of $\phi = 0.5$ were used, since melt conditions refer to a scaling of $R_g \propto N^\nu$ with exponent $\nu = 1/2$ that we expect for our plasmids at 0.05 M salt concentration. For this particular case, the results of Ref. [5] provide an estimate for the number average number of concatenations per ring,

$$f_n \approx \beta\phi R_g^2 \approx \beta\phi N. \tag{S4}$$

Here, $\beta$ is the numerical coefficient used to fit the data. We expect that the coefficient is related to the entanglement degree of polymerization in the melt, $\beta \propto N_e^{-1}$. For the constant $\phi$ of the simulations, the chain concentration is $\propto N^{-1}$ and thus, $P \propto N^{1/2}$. Since $f_n \propto N$ at constant $\phi$, $f_n$ is $\propto P^2$ for the large $P$ in Figure S5 (black solid line). This latter relation is used to extrapolate the data at large $P$ towards small $P$ for minimizing corrections due to the comparatively bulky monomers [15]. Gelation is expected for $f_n \approx 1$ [6] in perfect Olympic gels, which is reached for an extrapolated $P \approx 0.36$, see Figure S5. This overlap number provides a first estimate of the lowest concentration at which Olympic gels can be formed. We have to stress that the structure of Olympic gels is fully described by a single variable, which is $f_n$ [6]. Thus, we

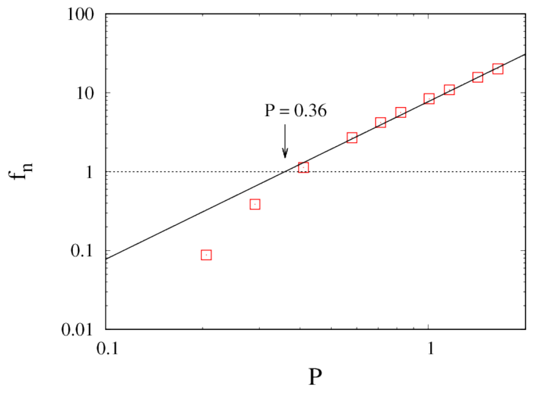


FIG. S5. Average number of concatenations per ring, $f_n$, as a function of the overlap number $P$ of concatenated rings at preparation conditions. Data taken from Ref. [5]. Black solid line is a fit $\propto P^2$ of the data around $P \approx 1$.

expect equivalent physical behavior of gels with the same $f_n$, as long as concatenated rings remain in the flexible limit, $N/f_n > 1$. This relation is key to compare between simulation data and experiments.

Let us apply the above estimate to the plasmid solutions. Our PVS10-DLK plasmid carries 11344 base pairs. Each base pair contributes on average 3.4Å to the length of a DNA strand. Thus, our plasmids have a contour length of approximately $L_c \approx 3.9\mu m$. The experiments are performed at a salt concentration of 0.05 M. At this salt concentration, a persistence length of $l_p \approx 55$nm was measured in Ref. [16] for the longest DNA fragment. The Kuhn length $b$ is approximately twice the persistence length for worm-like chains [9], $b \approx 2l_p$. The number of Kuhn segments, $N$, is $N \approx L_c/b$, which leads to $N \approx 35$ for our plasmids.

With these parameters, we find that the cubic gyration volume, $R_g^3$ of cyclic plasmids is approximately

$$R_g^3 \approx \left(\frac{b^2 N}{12}\right)^{3/2} \approx 5b^3. \tag{S5}$$

The bare diameter of a DNA double strand is $d \approx 2$nm and the volume of a Kuhn segment is $\approx \pi \left(d/2\right)^2 b$. Thus, a plasmid with $N \approx 35$ Kuhn segments of size $b$ occupies a bare volume of roughly

$$V \approx N\pi \left(d/2\right)^2 b \approx b^3/110. \tag{S6}$$

The density of double stranded DNA [17] is around $\rho \approx 1.7$ g/cm$^3$. Thus, a volume fraction of $V/R_g^3$ refers to a weight fraction of $\rho V/R_g^3 \approx \frac{1}{320}$g/ml within a volume of $R_g^3$. This weight fraction can be associated with an overlap number of $P = 1$. According to Figure S5, the gelation threshold is expected at an overlap number of 0.36. Therefore, we expect that Olympic gels may form above 0.11 wt% of plasmids in the solution,

$$c_{\text{gel}} \approx 0.36\rho V/R_g^3 \approx 1.1 \text{ mg/ml}. \tag{S7}$$

All experiments targeting Olympic gels were performed at weight fractions exceeding this threshold. Based upon the above estimate and assuming ideal Olympic gel structure, we expect that the samples at 0.125, 0.25, 0.5 and 0.8 wt% develop about 1.1, 2.2, 4.4, and 7.2 concatenations per plasmid. According to Ref. [6], the sample at 0.125 wt% should be very close to the gelation threshold.

Below, we test the above estimates by comparing different sets of experimental and simulation data. Note that for the shear simulations, we used the samples with 128, 256, 512, and 1024 monomers per ring that contain in average 2.7, 5.6, 10.9, and 20.2 pairwise concatenations per ring [18], respectively.

### S3. GEL ELECTROPHORESIS

Figure S6 summarizes all native agarose gel electrophoresis data. The control samples are predominantly a mixture of ≈ 91% of 11kb relaxed circular and about 9% of 11kb supercoiled DNA. Only at the largest concentration (0.25 wt%), a small amount of the DNA remains in the well.

As expected, supercoiled species are practically absent in the Olympic plasmids samples. Here, the gel electrophoresis data at the lowest concentration (0.01 wt%) show that the samples consist predominantly of relaxed circular DNA. The amount of linear DNA is about 20% for all concentrations, suggesting that a part of the rings opened, possibly due to the pipetting-induced shear forces during sample preparation or gel loading. Quantification of the linear DNA band density at higher concentrations is affected by the difficulty to accurately pipette the more viscous solutions. For all but the lowest concentration, a significant amount of DNA remains in the well, which is consistent with the formation of clusters of concatenated DNA with increasing concentration. The data of Ref. [6] show that the weight fraction of soluble species is dominated by single non-concatenated rings around and below the gelation concentration. We observe no clear bands of pairs or triplets of concatenated cycles in the data indicating that clusters of cyclic DNA may not penetrate the agarose gel in sufficient numbers for visualization, or that electrophoretic shear forces occasionally cause transient ring opening during the two hours of the gel run. Altogether, the presence of a growing immobile portion is a clear indication of structural changes inside the sample with increasing weight fractions of DNA.

### S4. RHEOLOGY

#### A. Solvent conditions, concentrations, and time scales

According to literature, charge screening of DNA is effective for concentrations exceeding 10 mM, see Figure 9 of Ref. [19] and the corresponding discussion in the text. Therefore, concepts for neutral polymers are suitable for discussing the rheology of our samples at 50 mM. Moreover, our buffer does not contain

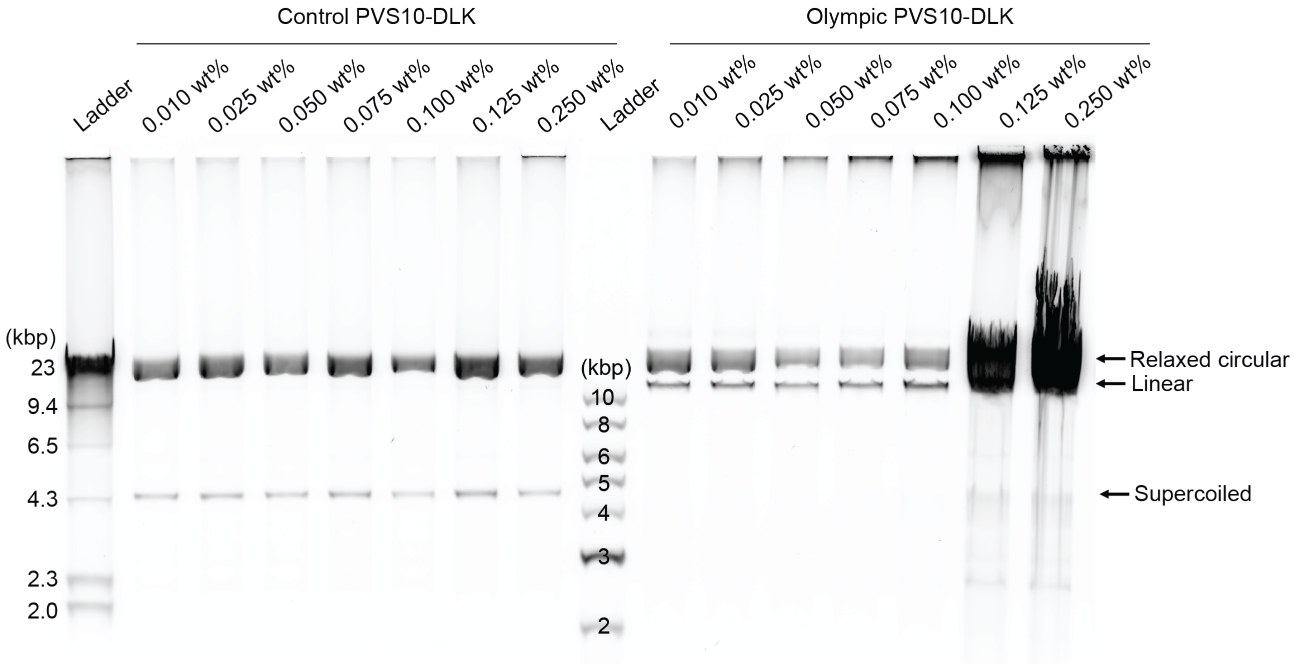


FIG. S6. Extended set of native agarose gel electrophoresis data of control and Olympic samples that were incubated at different concentrations. All samples were diluted to the same concentration immediately before gel loading. We note that samples containing high concentrations of Olympic gel PVS10-DLK were viscous and difficult to pipette, thus the dilution step possibly led to some mechanical disintegration of concatenated plasmid clusters.

multivalent salts. Therefore, we expect no effective attraction between DNA strands.

The relevant concentration scales for understanding the rheology of the DNA solutions are the transitions between the dilute and the semi-dilute regime at an overlap concentration $c^*$ and the transition between the semi-dilute non-entangled to the semi-dilute entangled regime at a concentration $c_{\rm e}$. According to Figure 2 of Ref. [19], these are observed for linear double stranded DNA at concentrations of $c^* \approx c^*_{\rm geo}/2$ and $c_{\rm e} \approx 3c^*_{\rm geo}$, respectively, where

$$c^*_{\rm geo} = \frac{3M}{4\pi R_g^3 N_A} \tag{S8}$$

is the geometric overlap concentration defined in Ref [19]. The average mass of a base pair is 618 Daltons, thus our plasmids have an average molar mass of approximately $7 \times 10^6$g/mol. With the $R_g^3$ of equation (S5), we obtain

$$c^*_{\rm geo} \approx 0.42 \text{mg/ml}. \tag{S9}$$

Thus, $c_{\rm gel} \approx 2.6c^*_{\rm geo}$, see equation (S7). Comparing with Ref. [19], we see that $c^*_{\rm geo}$ of our cyclic plasmids is about twice the $c^*_{\rm geo}$ of linear DNA with similar spatial extension (e.g. the 5.9 kbp sample in Table II of Ref. [19]). Entanglements arise from the local constraints imposed by neighboring molecules. These constraints are in first approximation a function of the total DNA concentration. Due to the larger $c^*_{\rm geo}$ of cyclic plasmids, we expect that the semi-dilute non-entangled regime is narrower when comparing with linear DNA due to a similar $c_{\rm e}$ for both kinds of samples. Thus, with respect to $c^*_{\rm geo}$ of cyclic DNA, we estimate that roughly $c_{\rm e} \approx 1.5c^*_{\rm geo}$. Therefore, all of our gelled samples are within the semi-dilute entangled regime.

Regarding the relevant time scales for dynamics, a rough orientation is obtained by considering the diffusion data of circular DNA in the dilute limit [20]. We focus on the circular DNA with 11.1 kbp that is almost identical to the 11.3 kbp of our plasmids. This circular DNA develops a diffusion coefficient of 1.31 $\mu$m$^2$/s. It was argued in literature [21, 22] that a scaling of $\nu = 0.5$ seems to be appropriate as we have used above. Therefore, we combine our estimate for $R_g \approx 0.188$ $\mu$m with the measured diffusion coefficient of the 11.1 kbp DNA to arrive at a rough estimate for the relaxation time of our cyclic DNA in the dilute limit, $\tau \approx R_g^2/D \approx 0.027$ s. Equilibrium properties of gels will arise on time scales clearly exceeding this lower bound.

### B. Olympic gels

For the relaxation time of the chains in the Olympic gel, we start with the approximation used for conventional networks: the relaxation time of the network is on the order of the Rouse time of the network strands (except for network defects as discussed below) [23]. The Rouse time of a linear chain in semi-dilute solutions is estimated as [9]

$$\tau \approx \frac{\eta_{\rm s} b^3}{kT} N^2 \phi^{(2-3\nu)/(3\nu-1)}, \tag{S10}$$

where $\eta_{\rm s}$ is the solvent viscosity, $k$ the Boltzman constant, and $T$ the absolute temperature. For $\nu = 0.5$, we obtain $\tau \propto \phi$ and thus, $\tau \propto (c/c^*)$ [9]. For the samples at the largest concentration with $c/c^* \approx 19$, we expect a similar growth of the Rouse relaxation time of linear plasmids to approximately 0.5 seconds, while the cyclic DNA will relax faster by a factor of $\approx 2$ due to the smaller $R_g$. The corresponding frequency of 2Hz (or 4 Hz for the cyclic DNA) is more than one order of magnitude above the time scale of the shallow $G''$ peak for a wt% of 0.8 in Figure 5g. Therefore, Rouse relaxation of linear chains or concatenated rings cannot explain the appearance of a $G''$ peak at a frequency around 0.1 Hz for 0.8 wt% DNA.

In conventional networks, slow relaxation modes are typically attributed to network defects like pending structures within the gel [24, 25]. However, these can be ignored for the sample at 0.8 wt% DNA: according to Figure 4 of Ref. [6], we expect only around 2% of network defects for an estimated $f_w \approx f_n + 1 \approx 8.2$. Therefore, network defects are not causing the $G''$ peak in Figure 5g.

According to the gel electrophoresis data, there is about 20% of linear DNA present inside the gel. For mixtures of linear and cyclic polymers, the last relaxation step of the cyclic polymers occurs when the penetrating linear chains relax [26]. In fact, ultra pure ring melts are necessary to remove the impact of the linear chains from rheological data on ring melts or solutions [26, 27]. For the reptation of entangled linear DNA inside the Olympic gel, we expect that the relaxation time scales roughly $\propto (c/c_e)^{7/3}$ for concentrations $c > c_e$. This provides an additional slow down by a factor of $(c/c_e)^{4/3} \approx 30$ for the sample at 0.8 wt% as compared to the Rouse time (based upon the above estimates for $c_{gel}$ and $c_e$). This leads to a relaxation of stress at frequencies around 0.1 Hz, in agreement with the location of the shallow $G''$ peak in Figure 5g. Therefore, this peak may originate from the relaxation of the entangled linear chains inside the gel. Equilibrium storage modulus data must be collected on times clearly exceeding the time scale of slowest relaxation given by the shallow $G''$ peak. This condition is met for the data at the lowest frequency analyzed.

#### C. Control samples

To create the primary control sample for our experiments, the PVS10-DLK plasmid was nicked at a single location using the Nb.BsmI nicking enzyme. This digestion step turned most supercoiled plasmids into relaxed plasmids to closely match the configuration of the Olympic plasmids while precluding mechanical interlocking. Notably, the relaxed plasmid control is more viscous than the original supercoiled plasmid solution at the same concentration. This finding mirrors previous studies reporting an unexpected elastic behavior for enzymatically digested plasmids as well as for mixtures of supercoiled plasmids with relaxed plasmids [28, 29]. This surprising behavior has been previously attributed to unusual entanglement processes or interpenetration that are suspected to introduce additional constraints [28–30]. Recall that 9% of the DNA in the control samples is still supercoiled, see section S3. In recent years, it was proposed that localized tip bubbles form under physiological conditions [31] at positions related to the AT rich domains along the supercoiled plasmid DNA [31–34]. If two tip bubbles come into contact, the exposed bases could optimize their free energy by partially associating with each other. In effect, it would be possible that a reversible gel of supercoils could be created entrapping the overlapping relaxed rings, providing an alternative explanation of the observations. Overall, even though the surprisingly high moduli of the control samples are not fully understood, they provide a reference for distinguishing the effects of permanent plasmid concatenations in the Olympic gel from transient effects in non-concatenated plasmid solutions.

#### D. Plateau modulus, network defects, and gelation threshold

As shown in Figure 4 of Ref. [6], the weight fraction of sol and of pending cycles (concatenated only once by the gel) disappears quickly with increasing number of concatenations. By comparison with these data, we expect that for the samples at the largest two concentrations, clearly more than 90% of the cycles are elastically active and thus, contribute to $G'$. In the limit of large $f_n \gg 1$, we expect that the scaling of the modulus becomes asymptotically the same as the plateau modulus of an entangled polymer solution [9], i.e. $G' \propto (c/c^*)^{7/3}$. This scaling is roughly confirmed by the data in Figure 5e. This is a much stronger dependence as expected for conventional gels at low concentrations where $G' \propto c$ in first order approximation [35].

For the transition region between the threshold concentration for gelation and the asymptotic limit at $f_n \gg 1$, the connectivity of the cyclic molecules matters. A constant fraction of linear chains in all samples plays no role for our discussion, if we compare data at different concentrations, where constant factors for all samples drop out. In a zero order approximation, we multiply the concentration $c$ with the weight fraction of elastically active rings, $w_{act}$, to estimate $G'$ according to $G' \propto (w_{act}c/c^*)^{7/3}$. Assuming that our estimates from section S2 for $f_n = 2.2$, 4.4, and 7.2 for the three largest concentrations are correct, we expect for the elastically active material from the computations of Ref. [6] that $w_{act} \approx 0.6$, 0.93, and 0.98 for three samples at the largest concentrations. Thus, we expect on a qualiatitive basis when comparing data of different samples that $G'(0.5)/G'(0.8) \approx (0.93/(1.6 \cdot 0.98))^{7/3} \approx 0.30$ and $G'(0.25)/G'(0.8) \approx (0.6/(3.2 \cdot 0.98))^{7/3} \approx 0.02$. The corresponding ratios of the experimental data are 0.26 and 0.03, respectively. Therefore, our estimate for $f_n$ appears to be not far from the experimental conditions despite the existence of a small portion of linear chains. Note that we have not included the lowest concentration at $f_n \approx 1.1$ in the discussion, since the mean-field estimate used to derive the theoretical prediction in Ref. [6] breaks down close to the gelation threshold at $f_n \approx 1$.

For a discussion of the absolute values of $G'$ let us consider the affine model for a "dry" network as reference case where a cyclic DNA with volume $V$ may contribute $kT$ to the plateau modulus. This results in

$$G'_{\mathrm{ref}} \approx \frac{kT}{V} \approx \frac{4.114 \times 10^{-21}\mathrm{J}}{b^3/110} \approx 3.4 \times 10^2 \mathrm{Pa}, \quad \text{(S11)}$$

A DNA content of 0.8 wt% refers to a volume fraction of 0.0047. Thus, $kT$ per DNA molecule provides a reference plateau modulus of $G'_{\mathrm{ref}} \approx 1.6$ Pa at 0.8 wt%, which is a factor of 4.6 below the measured plateau modulus of 11.9 Pa. In order to convert this factor into a rough estimate for $f_{\mathrm{n}}$, we have to recall that $f_{\mathrm{n}}$ is defined as the average number of rings that are entrapped by a given ring [5]. Depending on the model for entanglements [11, 36–40], an additional coefficient in between 1/3 and 1 emerges for the contribution of such an entanglement to the plateau modulus. For instance, for the slip-tube model [11] (as one of the most advanced models for rubber elasticity), a contribution of 4/7 kT per entangled chain section reflects the sliding of the entanglements upon deformation. With this estimate, we arrive at $f_{\mathrm{n}} \approx 8.1$, which is just about 13% larger than the estimate of $f_{\mathrm{n}} \approx 7.2$ based upon the concatenation data of the simulations. Correspondingly, the $G'$ data at 0.8 wt% provide an estimate for the gelation threshold around 0.1 wt% of plasmid DNA. This is in excellent agreement with our experimental data and the gelation discussion based upon the simulation data, see section S2.

#### E. Flow curves, amplitude sweeps and step strain data

Figure S7 shows the flow curves of the Olympic gels and the control samples. For this experiment, the shear rate was incresed exponentially as a function of time. Thus, the shear strain is roughly proportional to the shear rate. Except for a small systematic shift at low strains (probably due to some inaccuracy for finding the exact zero stress condition), the control samples develop a highly reproducible behavior between first and second measurement. Thus, all processes involved in the flow behavior equilibrate within the one minute break between the experiments. At high concentrations, we observe the formation of a stress peak followed by some apparent slippage regime at largest strain rates. The Olympic gels develop a similar stress peak at clearly higher stress and strain followed by slippage. In contrast to the control samples, this stress peak is largely modified in the repeat example. In general, the observed behaviour could result from either sample breakage or from wall slip of the samples in both cases. However, for identical concentrations, wall slip should occur for roughly the same forces acting on the DNA strands. Due to the clearly larger critical stress for the Olympic gels in the first run, wall slip can be excluded, and the samples start breaking at some critical strain. For the control samples, the connections across the broken interface are being rearranged roughly within the one minute break between the experiments. For the Olympic gels, it appears that only a small portion of the connections was re-established. This qualitative

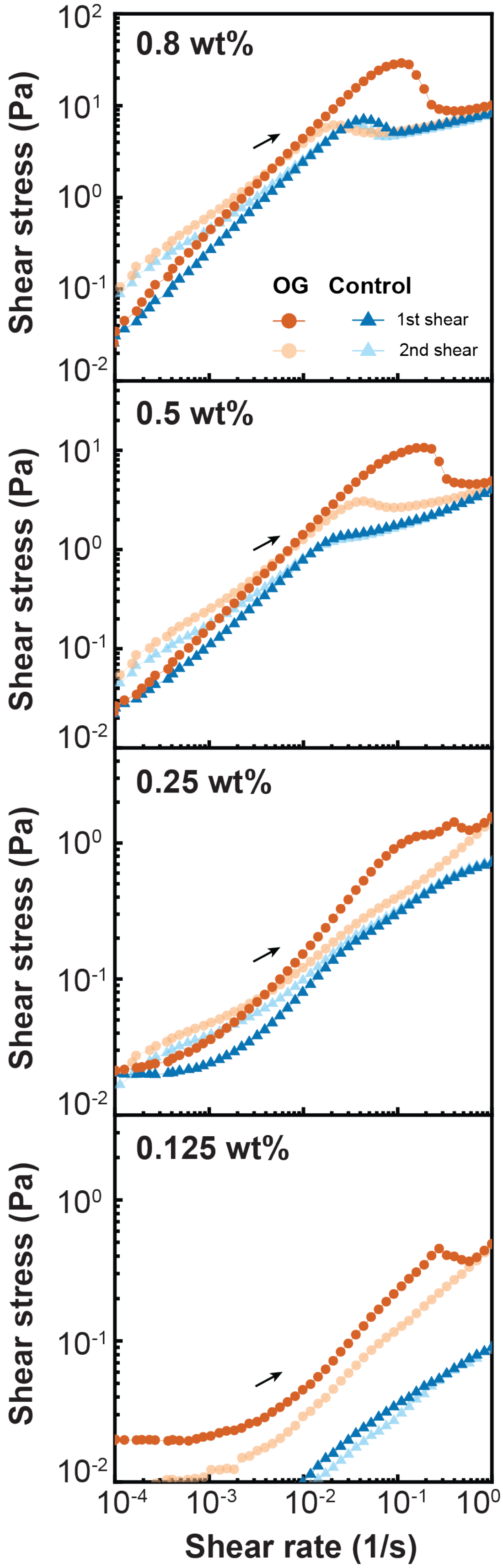


FIG. S7. Flow curves of Olympic gels and control. The samples were sheared rotationally at a shear rate from 0.0001 to 1 sec$^{-1}$ at 20°C for 2 consecutive repeats with a 1-minute break in between where the shear strain was reset to the point when the sample was under zero shear stress.

difference agrees well with the high predicted binding energy of the lock & key domains (Olympic gels): After breakage, the broken halfs of the samples slide with respect to each other. Newly formed concatenations across the broken interface carry a high load and will be opened again. Newly formed concatenations not across the broken interface will not be deformed and will be stable up to the livetime of lock & key domain. Thus, a continued slippage over a time span comparable or longer than the closing time of the rings will largely bias concatenation within one broken half of the sample. This bias will survive up to the livetime of the lock & key domain. In consequence, the sample breaks at a clearly lower stress in the repeat experiment. Besides this mechanism, it is possible that some DNA simply breaks or that finding the perfect position between lock & key domains may require significantly more time than the relaxation time of a DNA molecule. This latter argument agrees with the linear chains found in the DNA electrophoresis and AFM images. However, we have to recall that pipetting the samples in these experiments might have had an impact.

In this section, we discuss amplitude sweeps measured at 1 Hz and step strain data. The frequency of the amplitude sweeps was chosen to be close (a factor of $\approx 4$ less) to the frequency associated with the Rouse time of the rings providing sufficient time for the rings to rearrange their conformations upon deformation. Additional measurements at 0.1 Hz developed a much larger hysteresis at high strains indicating that some rings might open and rearrange their concatenations. The chosen frequency, thus, is a compromise for maximising the accessible strains without allowing for significant structural rearrangement of the gels during the time of the measurement.

In Figure S8, we show hysteresis measurements from two physical replicates for each sample where we averaged data of runs with increasing and decreasing strain. The low error bars for $G'$ demonstrate that our results are highly reproducible except for the Olympic gel at the lowest concentration, which is close to the gelation concentration. In our discussion below, therefore, we focus on the three largest concentrations.

In Figure S8, all control samples develop a pronounced shear thinning and flow behavior around a shear strain of roughly 100%. The loss modulus starts to dominate over the storage modulus at large strains. The amplitude sweep is fully reversible for all concentrations and develops no hysteresis similar to the flow data.

In Figure S8, the amplitude sweeps of the Olympic gels show a similar behavior as the control samples except for some subtle differences: as expected from the flow curves, there is a broader hysteresis between the up and down cycle due to a partial breakage of the sample. For oscillatory shear, a part of the reconnection occurs at low strains. Thus, the reconnection process is less biased as compared to the flow experiment leading to a gradual breakage of the gel that might occur in parallel to significant sliding of the rings. Ring opening and sample breakage becomes dominant at the intersection of $G'$ and $G''$. At strains below this intersection, the deformation behavior of the Olympic gel may stand out and is analyzed in Figure S9. On a quantitative level, shear strains in the range of 100% refer to an elongation of the molecules by about 40%. For flexible polymers, such a deformation roughly doubles the elastic energy per elastic strand providing $kT$ for sliding motion per entangled strand. Thus, a rearrangement of the ring conformations due to deformation should become effective at these strains. For these strains, there is also an excess of $kT$ of tensile energy along the ring contour that will not have much impact on the opening of the rings, since it remains clearly below the $\approx 40kT$ binding energy of the lock & key domain. For the largest three wt%, $G''$ exceeds $G'$ for strains in the range of 600% or longer. Here, due to the much higher elastic energy per ring at these strains, the rings start to open significantly and the data are no longer characteristic of a permanent Olympic gel. Above, we estimated $f_\text{n} \approx 7.2$ for the 0.8 wt% sample, which is between the $f_\text{n}$ measured for the $N = 256$ and $N = 512$ simulation samples, see section S2. An enlarged slippage at low strains was determined only for the samples with $N \geq 512$, i.e. for $f_\text{n} \gtrsim 10$, therefore, this feature must not show up yet in our rheology data. In Figure S9, we show the same measurement as in Figure S8, where we plot now stress vs. strain. At low strains, an almost perfectly linear stress-strain relation is observed. When comparing with the simulations in Figure S2, the corresponding strains of order 10% or below are close to the noise limit of the simulations. For strains exceeding 50%, a sublinear stress-strain relation is observed characterized by an effective exponent that changes slowly with increasing wt% and thus, $f_\text{n}$. For the three sets of data in Figure S9, we obtain effective exponents of $0.52 \pm 0.04$, $0.61 \pm 0.03$, and $0.65 \pm 0.01$ for increasing wt%. The transition point between both regimes moves to lower strains with increasing wt%. In the simulations with an average larger $f_\text{n}$ and polymer concentrations, we obtained $0.78 \pm 0.03$ averaged over the three samples with the largest $N$. Thus, the simulation data extend the trend observed in the experiment but might indicate also a partial loss of concatenations during the experiment by minor ring opening at the corresponding strains. This is supported by control experiments at lower frequency that lead to smaller effective exponents at high strains. This comparison motivated also the choice of a rather high frequency to minimize ring opening during the amplitude sweeps. Moreover, all other experiments (except for flow) were performed at low strains to suppress strain induced ring opening. The clearly sub-linear stress-strain relation of the Olympic gel samples in Figure S9 is a strong indication that the elasticity is dominated by entanglements. Altogether, the rheology data agree qualitatively with the computer simulations and support strongly that we have successfully synthesized Olympic gels.

The step strain data of the control samples at the largest wt% in Figure S10 shows a slow logarithmic decay of stress as a function of time that is mainly complete (about 95%) after about 1000 seconds. In contrast to the control samples, the Olympic gel samples

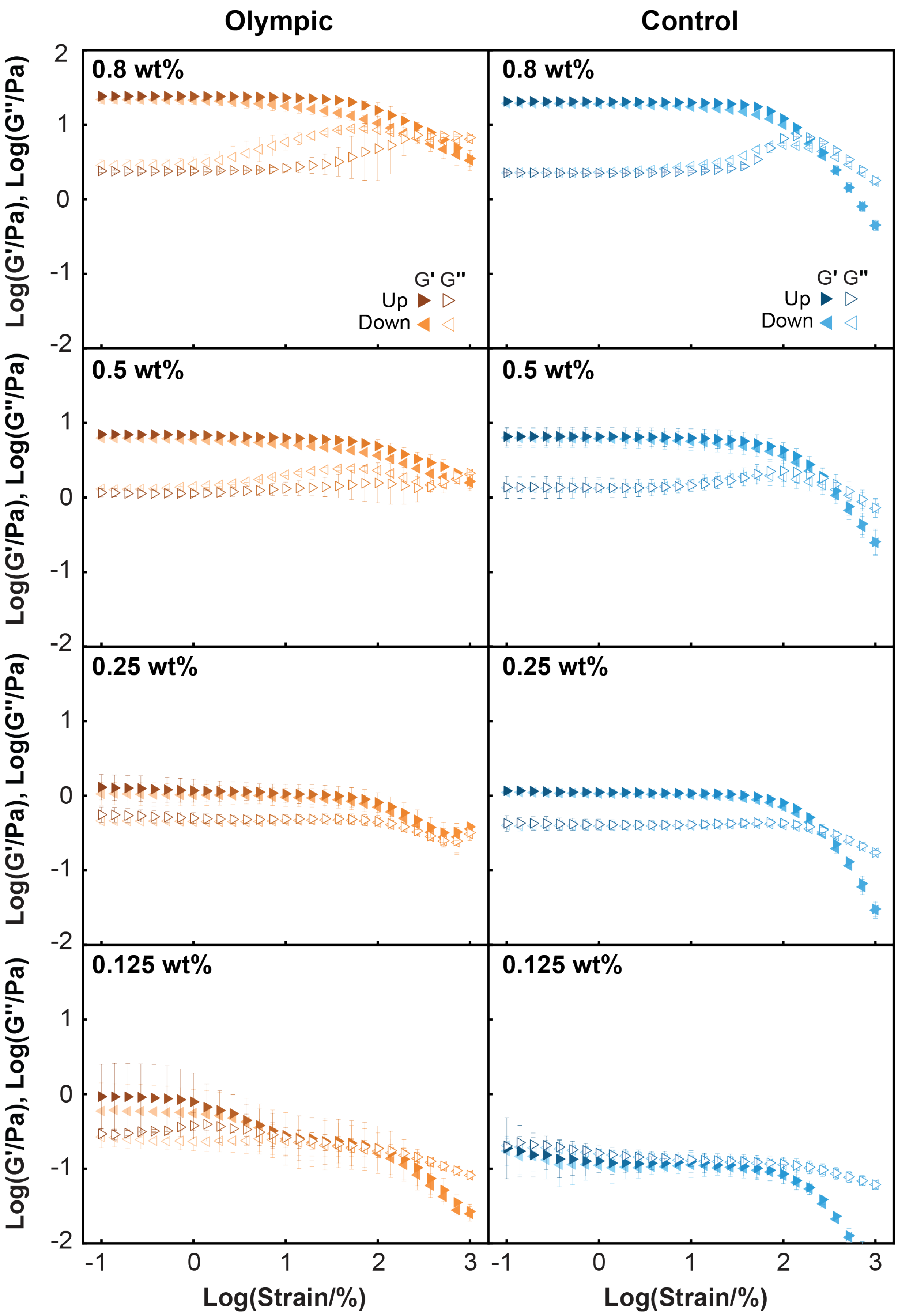


FIG. S8. Amplitude sweeps of Olympic gel and control. The first "up" sweep measured the sample from 0.1% to 1000% strain, followed by the second "down" sweep from 1000% to 0.1% strain. Both sweeps were performed at 1 Hz at 20°C. Data are shown as the mean ± s.d. (n = 2 physical replicates).

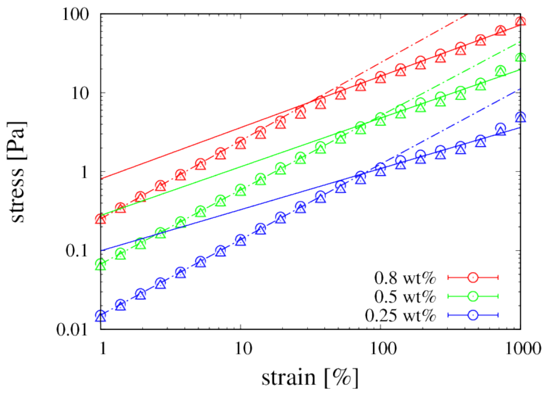


FIG. S9. Stress-strain curve of Olympic gels computed from the data of Figure S8 measured at 1Hz. Circles refer to a measurement for increasing strain, triangles refer to decreasing strain.

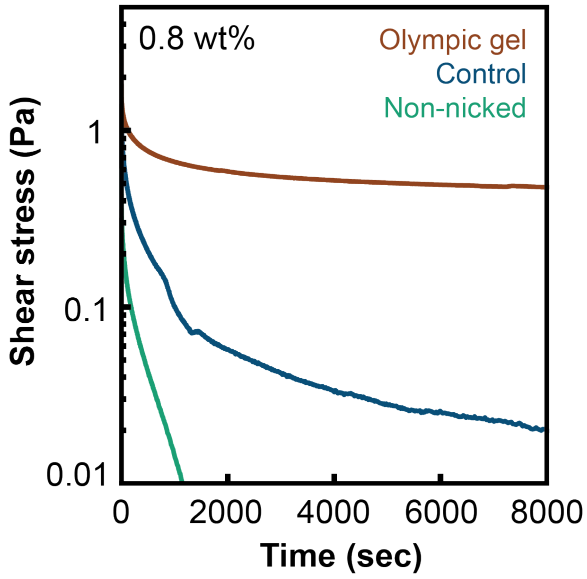


FIG. S10. Stress-relaxation curves of 0.8 wt% samples at 20 °C. The shear stress was recorded over time at a fixed strain of 15%.

may approach a plateau indicating that these samples behave like permanent gels on the time scale of the experiments.

### S5. SWELLING

In preceding work [41], it was shown that Olympic gels can develop an unexpected swelling behavior due to the existence of a desinterspersion process during swelling. Desinterspersion is driven by the pairwise repulsion of overlapping non-concatenated rings, which is a reasonable approximation for a low average number of concatenations and excluded volume interactions between the ring polymers. For the present set of data, however, chain conformations are close to ideal [21, 22] and excluded volume interactions are too weak for a mutual repulsion between pairs of rings.

Swelling equilibrium refers to the balance of the osmotic pressure with the “elastic pressure” of the network. For the former, one considers typically a virial expansion [9] of the form

$$\Pi = \frac{kT}{V_\mathrm{b}}\left[\frac{\phi}{N} + \frac{\mathrm{v}\phi^2}{2V_\mathrm{b}} + \frac{\mathrm{w}\phi^3}{3V_\mathrm{b}^2} + ...\right]. \qquad \text{(S12)}$$

Here $V_\mathrm{b} = \pi\,(d/2)^2\,b$ is the bare volume of a Kuhn segment, whereas v and w are the excluded volume parameter and the three-body interaction parameter, respectively. For our systems, an “effective width” (due to electrostatic interactions) of the DNA of roughly $d_\mathrm{eff} \approx 10$ nm [42] and a Kuhn length of $b \approx 110$ nm [16] have been measured. For semi-flexible polymers, there is $\mathrm{v} \approx b^2 d_\mathrm{eff}$ and $\mathrm{w} \approx b^3 d_\mathrm{eff}^3$ [9]. Moreover, in our study, all volume fractions are $\phi \lesssim 0.01$ and there is $N \approx 35$. Thus, we arrive at the unusual situation that even though the chain conformations are ideal, $\mathrm{v} \approx 0$, osmotic pressure is still dominated by two body interactions. Moreover, since there is less than $kT$ of osmotic pressure per ring (and thus, less than $kT$ repulsion between the rings), we do not expect that the push-off regime for $P \approx 1$ discussed in Ref. [41] contributes significantly to the swelling equilibrium. Instead, we expect the same scaling as expected for conventional polymer networks in the entangled limit. Here, it has been proposed that $Q \propto \phi_0^{-1}$ for swelling in good solvents, see equation (S17) of Ref. [43], which we assume due to the dominating two body interactinos. This implies that $\phi/\phi_0$ is constant at swelling equilibrium. The samples with the largest wt% of DNA seem to approach this trend, see Figure 5c. Note that this trend is largely different from conventional gels in the non-entangled limit with an expected $Q \propto \phi_0^{-1/4}$ [9], which would lead to a significantly decaying ratio of $\phi/\phi_0 \propto \phi_0^{-3/4}$ and an increasing volume ratio $V/V_0 \propto \phi_0^{3/4}$, where $V$ and $V_0$ are the sample volume at preparation and at swelling equilibrium, respectively.

In Figure 5b, all Olympic gels develop a maximium degree of swelling before the concentration decays. Recall that the Olympic gels contain some linear DNA that can diffuse out of the gel. This may cause a slow decay of the gel volume after the initial rapid swelling driven by the solvent diffusion. The observed rather sharp concentration dependent transition between dissolution and swelling (Figure 5c) is to be expected, since theoretically, the equilibrium degree of swelling diverges at the gelation threshold. Note that this divergence must be followed by a decay towards larger wt% before crossing over to the ultimate scaling at large wt%. This process combined with a slow growth of $V/V_0 \propto \phi_0^{1/4}$ as predicted by Ref. [41], for instance, could provide an alternate explanation of the data in Figure 5c. Therefore, a final conclusion about the scaling of the swelling Olympic gels remains difficult. However, all observations related to equilibrium swelling support the existence of a gelation concentration around 0.1 wt% DNA and the formation of gels that withstand dissolution over at least one week.

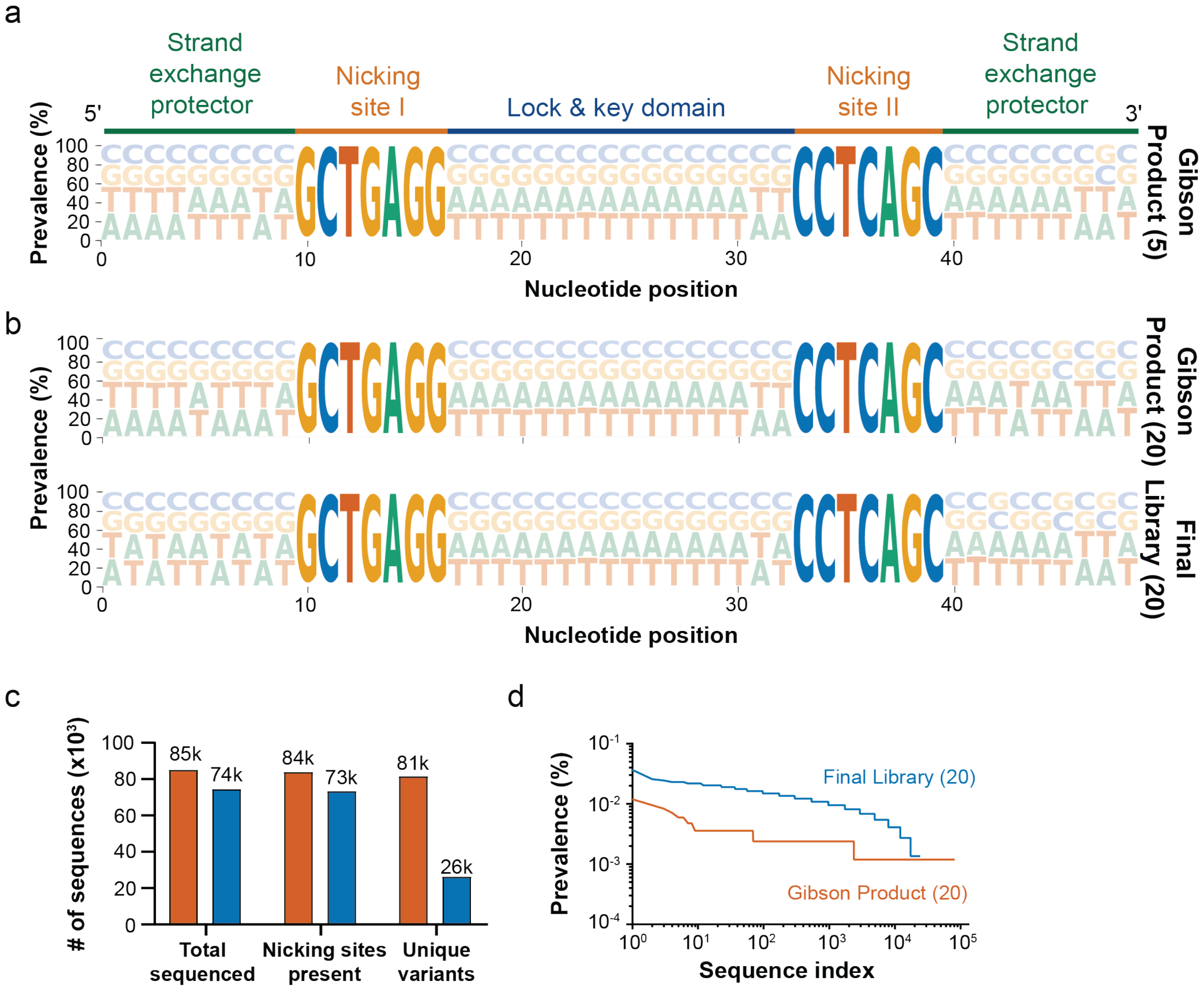


FIG. S11. a) Comparison of the prevalence of the four canonical bases within the DLK insert in the Gibson product before transformation into E. coli when 5 parallel transformations were combined for large-scale growth of the PVS10-DLK library. b) Comparison of the prevalence of the four canonical bases within the DLK insert in the Gibson product before transformation into E. coli and in the final library when 20 parallel transformations were combined for large-scale growth of the PVS10-DLK library. The size of each letter corresponds to the probability of the indicated nucleotide to appear at that position. c) Next-generation sequencing data for the final library using 20 transformations. Orange bars indicate the sequences from the Gibson product before transformation, and blue bars indicate the sequences from the final PVS10-DLK library. d) Comparison of the prevalence between sequence variants (with nicking sites present) in descending order for the final library using 20 transformations.

### S6. DISSOLUTION

Dissolution of a polymer solution or a reversible gel of DNA plamids is a complex phenomenon involving several physical processes that might affect the time dependence of this process [44]. In our particular case, we can simplify the discussion by considering that the relaxation time of the molecules (seconds) is orders of magnitude below the time scale of dissolution (hours to days). Therefore, relaxation phenomena in the surface layer or inside the gel play no essential role for the dissolution time and we are left with considering diffusion of solvent, detached DNA, or the time scale for detaching the DNA from a reversible gel as rate limiting steps for dissolution.

The fastest of these processes is the diffusion of the solvent that causes the observed swelling of the gel layer at short times, see Figure 5b. Following the rapid apparent volume increase of the plasmid-rich solution by diffusion of solvent, the control plasmids then slowly dissipate.

In order to understand whether this disintegration or the diffusion of the DNA out of the layer are the time limiting steps, we consider first the diffusion time of the plasmids. Based upon the diffusion data of Ref. [20] and the discussion of section S4 A, we estimate a diffusion time of about 53 hours for our plasmids for a distance of 500 $\mu$m. This is slightly longer than the time scale at which the DNA of the control samples disappears from the bottom layer shown in Figure 5a. Therefore, the lifetime of any reversible bond between the plasmids must be shorter than the timescale of plasmid diffusion.

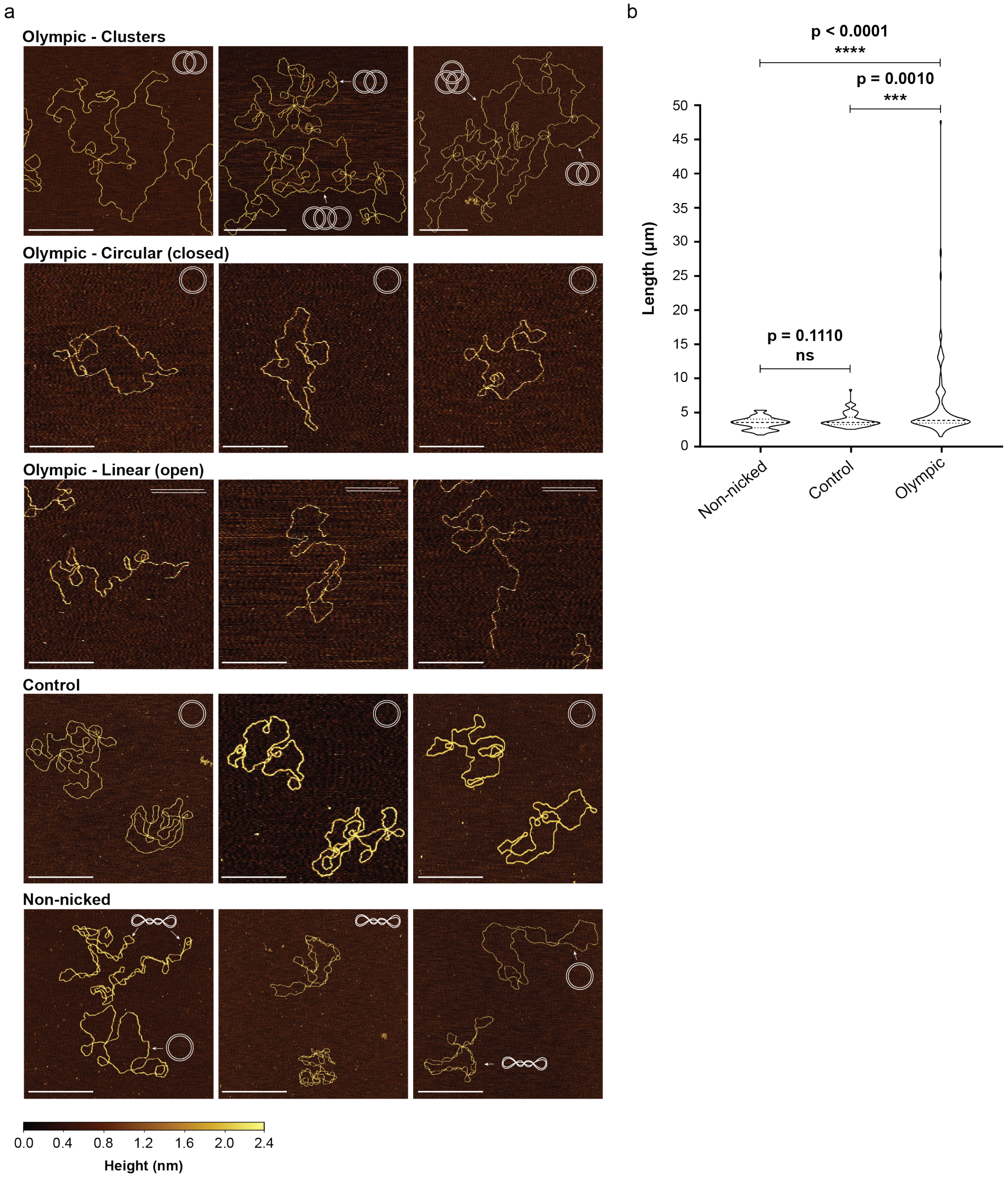


FIG. S12. a) Representative examples of different plasmid forms seen in AFM imaging for Olympic, control, and non-nicked samples diluted to 0.0005 wt% after annealing. The white scale bar is equal to 500 nm. The color scale bar indicates height in the z-direction. b) Contour lengths of plasmid objects for each sample ($n = 67$ for non-nicked; $n = 74$ for control; $n = 103$ for Olympic. A non-parametric Mann-Whitney U test was used to determine statistical significance between samples).

**0.25 wt% Olympic gel**

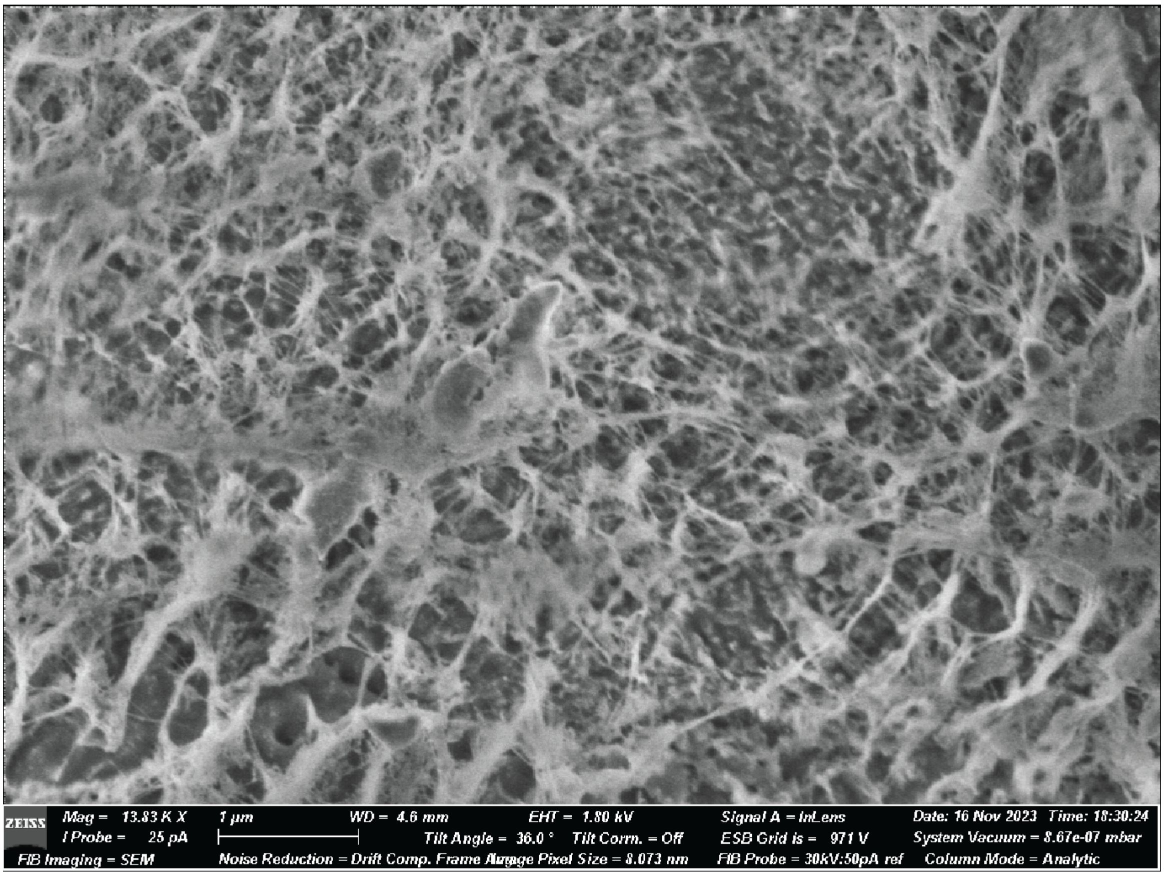


**0.25 wt% Control**

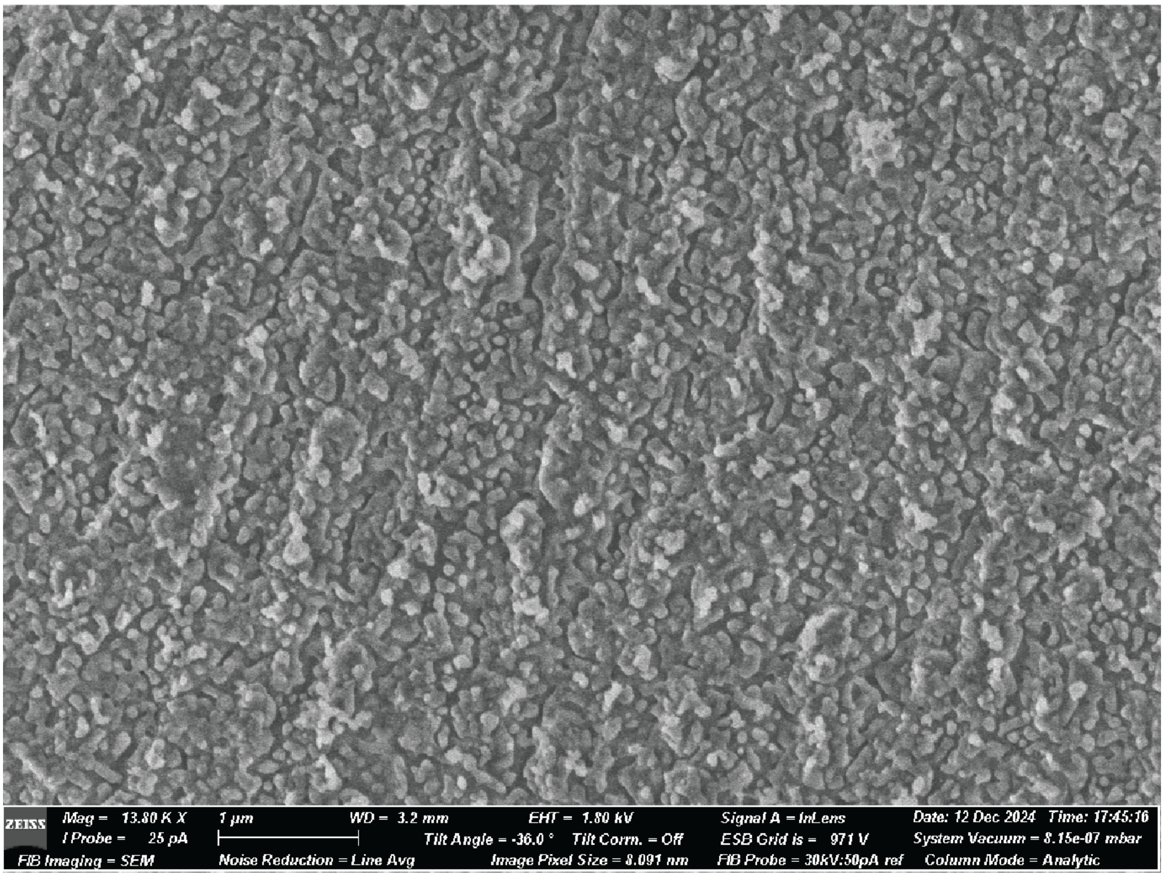


FIG. S13. Uncropped cryo-SEM images for Olympic gel and relaxed control samples annealed at 0.25 wt% with accompanying instrument parameters.

### 0.25 wt% Non-nicked

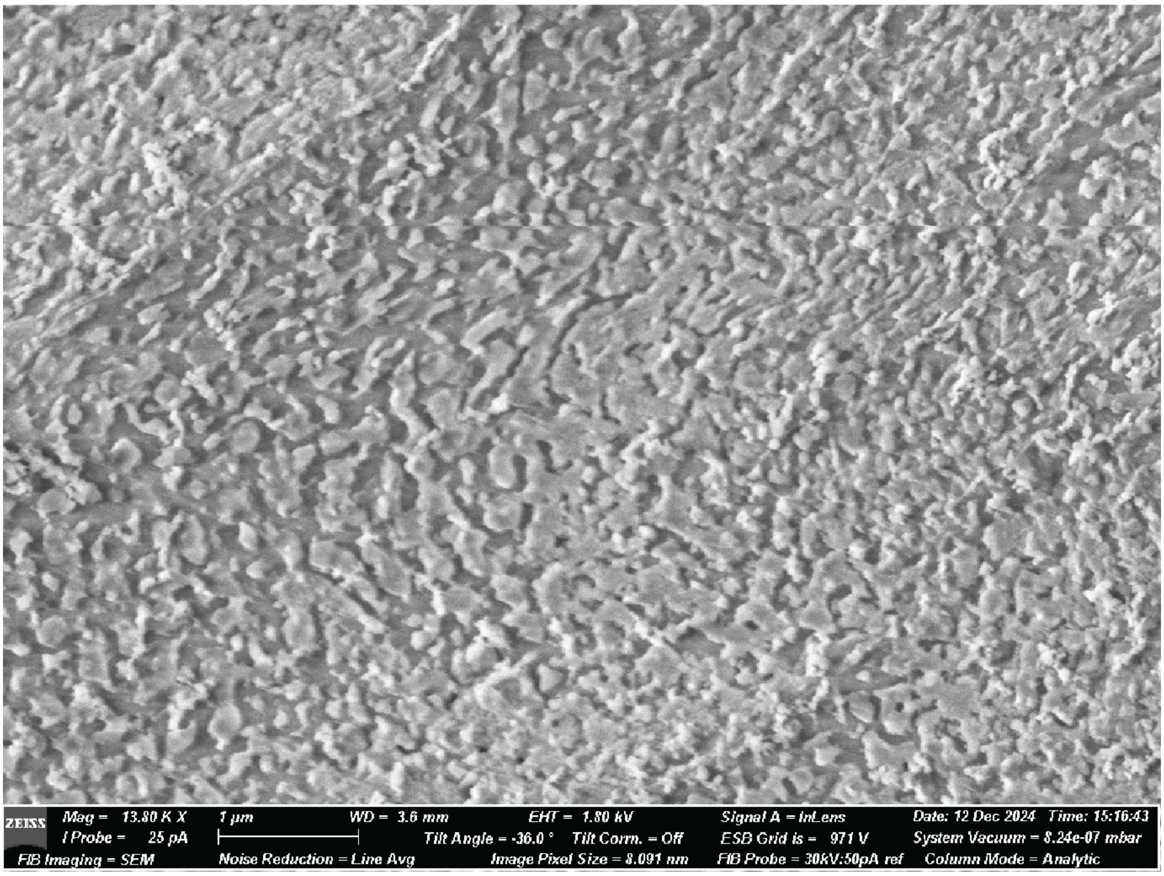


FIG. S14. Uncropped cryo-SEM images for non-nicked samples annealed at 0.25 wt% with accompanying instrument parameters.

---


[1] I. Carmesin and K. Kremer, Macromolecules **21**, 2819 (1988).

[2] H. P. Deutsch and K. Binder, J. Chem. Phys. **94**, 2294 (1991).

[3] T. Müller, M. Wengenmayr, R. Dockhorn, H. Rabbel, M. Knespel, A. Checkervarty, V. Sinapius, Y. Guo, and M. Werner, “Lemonade-project/lemonade: Lemonade v2.2.2,” (2021).

[4] T. Müller, M. Wengenmayr, R. Dockhorn, and M. Knespel, “Lemonade-project/lemonade-gpu: Release v1.2,” (2021).

[5] M. Lang, J. Fischer, and J.-U. Sommer, Macromolecules **45**, 7642 (2012).

[6] J. Fischer, M. Lang, and J. U. Sommer, Journal of Chemical Physics **143**, 243115 (2015).

[7] T. Müller, J.-U. Sommer, and M. Lang, Macromolecules **55**, 7540 (2022).

[8] M. Lang and T. Müller, Macromolecules **55**, 8950 (2022).

[9] M. Rubinstein and R. H. Colby, *Polymer Physics* (Oxford University Press, 2003).

[10] P. M. Rauscher, K. S. Schweizer, S. J. Rowan, and J. J. de Pablo, Macromolecules **53**, 3390 (2020).

[11] M. Rubinstein and S. Panyukov, Macromolecules **35**, 6670 (2002).

[12] R. C. Ball, M. Doi, S. F. Edwards, and M. Warner, Polymer **22**, 1010 (1981).

[13] S. F. Edwards and T. A. Vilgis, Polymer **27**, 483 (1986).

[14] G. Heinrich, E. Straube, and G. Helmis, Adv. Pol. Sci. **85**, 33 (1987).

[15] M. Lang, Macromolecules **46**, 1158 (2013).

[16] A. Brunet, C. Tardin, L. Salome, N. Rousseau, P. Destainville, and M. Manghi, Macromolecules **48**, 3641 (2015).

[17] F. E. Arrighi, M. Mandel, J. Bergendahl, and T. C. Hsu, Biochemical Genetics **4**, 367 (1970).

[18] In Table I of Ref. [41], the $f_{\mathrm{n}}$ value of sample #7 with $N = 1024$ was confused with another sample ($N = 764$) of Ref. [5]. The present work and all plots in Refs. [5, 41] contain the correct $f_{\mathrm{n}} = 20.2$.

[19] S. Pan, D. A. Nguyen, P. Sridhar, and J. R. Sunthar, P. Prakash, J. Rheol. **58**, 339 (2014).

[20] R. M. Robertson, S. Laib, and D. E. Smith, PNAS **103**, 7310 (2006).

[21] D. R. Tree, A. Muralidhar, P. S. Doyle, and K. D. Dorfman, Macromolecules **46**, 8369 (2013).

[22] M. L. Mansfield, A. Tsortos, and J. F. Douglas, J. Chem. Phys. **143**, 124903 (2015).

[23] A. A. Gurtovenko and A. Blumen, Adv. Polym. Sci. **182**, 171 (2005).

[24] J. G. Curro and P. Pincus, Macromolecules **16**, 559 (1983).

[25] M. Lang, D. Göritz, and S. Kreitmeier, *Constitutive Models for Rubber IV*, edited by P. Austrell and L. Kari (Taylor & Francis Group, plc, London, UK,

2005) pp. 349–359.
[26] M. Kapnistos, M. Lang, D. Vlassopoulos, W. Pyckhout-Hintzen, D. Richter, D. Cho, T. Chang, and M. Rubinstein, Nature Materials **7**, 997 (2008).
[27] Y. Doi, K. Matsubara, Y. Ohta, T. Nakano, D. Kawaguchi, Y. Takahashi, A. Takano, and Y. Matsushita, Macromolecules **48**, 3140 (2015).
[28] K. R. Peddireddy, M. Lee, Y. Zhou, S. Adalbert, S. Anderson, C. M. Schroeder, and R. M. Robertson-Anderson, Soft Matter **16**, 152 (2020).
[29] D. Michieletto, P. Neill, S. Weir, D. Evans, N. Christ, V. A. Martinez, and R. M. Robertson-Anderson, Nature Communicationss **13**, 4389 (2022).
[30] W. A. Paiva, S. D. Alakwe, J. Marfai, M. V. Jennison-Henderson, R. A. Achong, T. Duche, A. A. Weeks, R.-A. R. M., and N. J. Oldenhuis, Adv. Mater. **36**, 2405490 (2024).
[31] C. Matek, T. E. Oulridge, J. P. K. Doye, and A. A. Louis, Scientific Reports **5**, 7655 (2015).
[32] J. S. Mitchell, C. A. Laughton, and S. A. Harris, Nucleic Acids Research **39**, 3928 (2011).
[33] S. H. Kim, M. Ganji, E. Kim, J. van der Torre, E. Abbondanzieri, and C. Dekker, eLive **7**, e36557 (2018).
[34] M. Burman and A. Noy, Phys. Rev. Lett. **134**, 038403 (2025).
[35] M. Lang, R. Scholz, L. Löser, C. Bunk, N. Fribiczer, S. Seiffert, F. Böhme, and K. Saalwächter, Macromolecules **55**, 5997 (2022).
[36] G. Heinrich and E. Straube, Acta Polymerica **34**, 589 (1983).
[37] G. Heinrich and E. Straube, Acta Polymerica **35**, 115 (1984).
[38] G. Heinrich and M. Kaliske, Comp. Theor. Pol. Sci. **7**, 227 (1997).
[39] R. G. Larson, T. Sridhar, L. G. Leal, G. H. McKinley, A. E. Likhtman, and T. C. B. McLeish, J. Rheol. **47**, 809 (2003).
[40] R. Everaers, Phys. Rev. E **86**, 022801 (2012).
[41] M. Lang, J. Fischer, M. Werner, and J.-U. Sommer, Physical Review Letters **112**, 1 (2014).
[42] E. G. Yarmola, M. I. Zarudnaya, and Y. S. Lazurkin, J. Biomol. Struct. Dyn. **2**, 981 (1985).
[43] T. Yamamoto, J. A. Campbell, S. Panyukov, and M. Rubinstein, Macromolecules **55**, 3588 (2022).
[44] B. A. Miller-Chou and J. A. Koenig, Prog. Polym. Sci. **28**, 1223 (2003).
